# The transcriptomic and spatial organization of telencephalic GABAergic neuronal types

**DOI:** 10.1101/2024.06.18.599583

**Authors:** Cindy T. J. van Velthoven, Yuan Gao, Michael Kunst, Changkyu Lee, Delissa McMillen, Anish Bhaswanth Chakka, Tamara Casper, Michael Clark, Rushil Chakrabarty, Scott Daniel, Tim Dolbeare, Rebecca Ferrer, Jessica Gloe, Jeff Goldy, Junitta Guzman, Carliana Halterman, Windy Ho, Mike Huang, Katelyn James, Beagan Nguy, Trangthanh Pham, Kara Ronellenfitch, Eric D. Thomas, Amy Torkelson, Chelsea M. Pagan, Lauren Kruse, Nick Dee, Lydia Ng, Jack Waters, Kimberly A. Smith, Bosiljka Tasic, Zizhen Yao, Hongkui Zeng

## Abstract

The telencephalon of the mammalian brain comprises multiple regions and circuit pathways that play adaptive and integrative roles in a variety of brain functions. There is a wide array of GABAergic neurons in the telencephalon; they play a multitude of circuit functions, and dysfunction of these neurons has been implicated in diverse brain disorders. In this study, we conducted a systematic and in-depth analysis of the transcriptomic and spatial organization of GABAergic neuronal types in all regions of the mouse telencephalon and their developmental origins. This was accomplished by utilizing 611,423 single-cell transcriptomes from the comprehensive and high-resolution transcriptomic and spatial cell type atlas for the adult whole mouse brain we have generated, supplemented with an additional single-cell RNA-sequencing dataset containing 99,438 high-quality single-cell transcriptomes collected from the pre- and postnatal developing mouse brain. We present a hierarchically organized adult telencephalic GABAergic neuronal cell type taxonomy of 7 classes, 52 subclasses, 284 supertypes, and 1,051 clusters, as well as a corresponding developmental taxonomy of 450 clusters across different ages. Detailed charting efforts reveal extraordinary complexity where relationships among cell types reflect both spatial locations and developmental origins. Transcriptomically and developmentally related cell types can often be found in distant and diverse brain regions indicating that long-distance migration and dispersion is a common characteristic of nearly all classes of telencephalic GABAergic neurons. Additionally, we find various spatial dimensions of both discrete and continuous variations among related cell types that are correlated with gene expression gradients. Lastly, we find that cortical, striatal and some pallidal GABAergic neurons undergo extensive postnatal diversification, whereas septal and most pallidal GABAergic neuronal types emerge simultaneously during the embryonic stage with limited postnatal diversification. Overall, the telencephalic GABAergic cell type taxonomy can serve as a foundational reference for molecular, structural and functional studies of cell types and circuits by the entire community.

## INTRODUCTION

The telencephalon, the most anterior part of the mammalian brain, comprises several large structures that are considered as the top-level command centers of the hierarchically organized brain networks and play integrative roles in information processing and generation of behavior and cognition. The telencephalon is composed of two major structures, cerebral cortex (as the shell) and cerebral nuclei (as the core), which arise from pallium and subpallium, respectively, of the developmental telencephalon. Cerebral cortex consists of isocortex, hippocampal formation, olfactory areas and cortical subplate, whereas the cerebral nuclei consist of striatum and pallidum. Within each of these major brain structures, there are multiple functionally specific regions and subregions (**Supplementary Table 1** provides the anatomical ontology from the Allen Mouse Brain Common Coordinate Framework version 3 (CCFv3)^1^ with full names and acronyms of all telencephalic regions), each comprising many cell types.

In the mouse cerebral cortex (CTX) (**Supplementary Table 1**), isocortex contains ∼35 cortical areas, including visual, auditory, somatosensory, gustatory, visceral and motor areas, as well as association areas in the prefrontal, medial and lateral parts. Hippocampal formation (HPF) is divided into hippocampus (HIP) and retrohippocampal regions (RHP), and the latter is further divided into medial and lateral entorhinal cortex (ENTm and ENTl), parasubiculum (PAR), postsubiculum (POST), presubiculum (PRE), subiculum (SUB), prosubiculum (ProS), hippocampo-amygdalar transition area (HATA) and area prostriata (APr). Olfactory areas (OLF) contain the entire olfactory sensory pathway, including main and accessary olfactory bulbs (MOB and AOB), anterior olfactory nucleus (AON), taenia tecta (TT), dorsal peduncular area (DP), piriform area (PIR), nucleus of the lateral olfactory tract (NLOT), cortical amygdalar area (COA), piriform-amygdalar area (PAA), and postpiriform transition area (TR). Cortical subplate (CTXsp) contains claustrum (CLA), endopiriform nucleus (EP), and lateral, basolateral, basomedial and posterior amygdalar nuclei (LA, BLA, BMA and PA).

In the mouse cerebral nuclei (CNU) (**Supplementary Table 1**), striatum (STR) consists of striatum dorsal region (STRd, also called caudoputamen, CP), striatum ventral region (STRv), lateral septal complex (LSX), and striatum-like amygdalar nuclei (sAMY). STRv is further divided into nucleus accumbens (ACB), fundus of striatum (FS), and olfactory tubercle (OT). LSX is further divided into lateral septal nucleus (LS), septofimbrial nucleus (SF) and septohippocampal nucleus (SH). And sAMY is further divided into anterior amygdalar area (AAA), bed nucleus of the accessory olfactory tract (BA), central amygdalar nucleus (CEA), intercalated amygdalar nucleus (IA), and medial amygdalar nucleus (MEA). Pallidum (PAL) consists of four subdivisions, with the dorsal region (PALd) containing globus pallidus external and internal segments (GPe and GPi), the ventral region (PALv) containing substantia innominata (SI) and magnocellular nucleus (MA), the medial region (PALm) containing medial septal nucleus (MS), diagonal band nucleus (NDB) and triangular nucleus of septum (TRS), and the caudal region (PALc) containing bed nuclei of the stria terminalis (BST) and bed nucleus of the anterior commissure (BAC).

From these highly complex regional subdivisions, a general organizing principle of the telencephalon with several parallel cortico-striato-pallidal circuit pathways has been revealed^2^. In a highly simplified view, the dorsal pathway from isocortex to CP to GPe/GPi mediates sensory/motor functions. On the ventral side, the prefrontal cortex-ACB-PALv and the LA/BLA-CEA-BST pathways mediate affective functions. The hippocampal-septal pathway along the medial axis mediates learning and cognitive functions. Underlying these complex circuit networks are an extraordinary array of neuronal cell types. Previous studies have revealed highly diverse and heterogeneous cellular properties of both glutamatergic and GABAergic neurons in the telencephalon, which likely contribute critically to the specific functions of different brain regions. To understand how the variety of brain functions emerge from this complex system, it is essential to gain comprehensive knowledge about the cell types, their regional/spatial specificity, and their inter-relatedness.

Glutamatergic excitatory neurons are the dominant neuronal class of the cerebral cortex and are generated within the ventricular and subventricular zones of the developing pallium. GABAergic inhibitory neurons are the dominant neuronal class of the cerebral nuclei and are generated in the ganglionic eminences of the developing subpallium. GABAergic neurons also migrate to the pallium and populate all parts of the CTX, intermingling with the glutamatergic neurons. In this study, we focus on telencephalic GABAergic neurons, which play a plethora of circuit functions. In CTX, they are mostly local inhibitory interneurons, modulating circuit dynamics, excitation-inhibition balance, and rhythmic activities. In CNU, they are mostly long-range inhibitory projection neurons and transmit circuit-specific information, though some are local interneurons as well and others (i.e., cholinergic neurons) play neuromodulatory roles.

The telencephalic GABAergic neuronal types arise mostly from the five principal progenitor domains of the subpallium: the medial ganglionic eminence (MGE), the caudal ganglionic eminence (CGE), the lateral ganglionic eminence (LGE), the embryonic septum, and the embryonic preoptic area (POA). Most GABAergic cell types are produced during the embryonic period and migrate along defined routes to disperse throughout the forebrain^3–5^. Cell fate specification in the progenitor domains is orchestrated by a combination of transcription factors and morphogens^4–13^.

Leveraging the comprehensive and high-resolution transcriptomic and spatial cell type atlas for the whole mouse brain we have generated^14^, supplemented with an additional scRNA-seq dataset collected from the pre- and postnatal developing brain, we conducted a systematic and in-depth analysis of the transcriptomic and spatial organization of GABAergic neuronal types in all regions of the mouse telencephalon and their developmental origins. We identified an extraordinarily large set of highly distinct cell types as well as continuous molecular gradients within and across different regions. These two aspects of the cell type landscape collectively shape the cellular diversity that underlie the diverse function of the many regions and neural circuits in the telencephalon. We discovered a comprehensive set of transcription factors (TF) that define all major subclasses and supertypes of the adult-stage telencephalic GABAergic neurons. We found strong expression of these TF marker genes in specific regions of the developing telencephalon, and thus were able to infer the developmental origins of all the GABAergic neuronal types described here, which had only been partially known before. The results reveal two prominent features of the telencephalic GABAergic neurons: 1) transcriptomically and developmentally related cell types are often found in far-apart and distinct brain regions, suggesting long-distance migration and dispersion is a common characteristic of nearly all classes of telencephalic GABAergic neurons; 2) cortical and striatal GABAergic neurons undergo extensive postnatal diversification, whereas septal and most pallidal GABAergic neuronal type repertoire emerges in an apparent burst in embryonic stage with limited postnatal diversification.

## RESULTS

### A transcriptomic and spatial atlas of GABAergic neuronal types in the mouse telencephalon

Previously, we reported the creation of a high-resolution transcriptomic and spatial cell type atlas covering the entire adult mouse brain based on the combination of single-cell RNA-sequencing (scRNA-seq) and spatially resolved transcriptomics using MERFISH (**Supplementary Table 2**)^14^. We defined a hierarchically organized whole-mouse-brain (WMB) cell type atlas comprising four nested levels of classification: 34 classes, 338 subclasses, 1,201 supertypes and 5,322 clusters. Neuronal cell types exhibit extraordinary diversity and constitute a large proportion of the whole brain cell type atlas, including 29 classes (85%), 315 subclasses (93%), 1,156 supertypes (96%) and 5,205 clusters (98%). However, in this previous study we only described the organization of neuronal cell types across the whole mouse brain at a coarse level (class – subclass). Here, we conduct a more in-depth analysis and introduce the most complete to-date taxonomy of all the GABAergic neuronal types in the entire telencephalon, at all levels of the hierarchy.

The telencephalic GABAergic neuronal cell type taxonomy, defined as the Subpallium-GABA neighborhood in the WMB cell atlas^14^, contains subclasses 39-90 that belong to 7 classes: OB-IMN GABA, CTX-CGE GABA, CTX-MGE GABA, CNU-MGE GABA, CNU-LGE GABA, LSX GABA, and CNU-HYa GABA (**Figure 1**). Contained within the total of 52 subclasses, there are 284 supertypes and 1,051 clusters, with a total of 611,423 high-quality single-cell transcriptomes (10x v2: 269,307 cells; 3,567 ± 1,264 genes per cell; 9,328 ± 5,502 UMIs per cell; 10x v3: 342,116 cells; 5,949 ± 1,625 genes per cell; 26,476 ± 14,943 UMIs per cell) (**Supplementary Table 3**). We provide several representations of this atlas for further analysis: a dendrogram at supertype resolution along with bar graphs displaying various metadata information (**Figure 1a**), UMAPs at single-cell resolution colored with different types of metadata information (**Figure 1b-d**), and constellation diagrams to depict multi-dimensional relationships among different subclasses and supertypes (**Figure 1e,f**).

**Figure 1.**
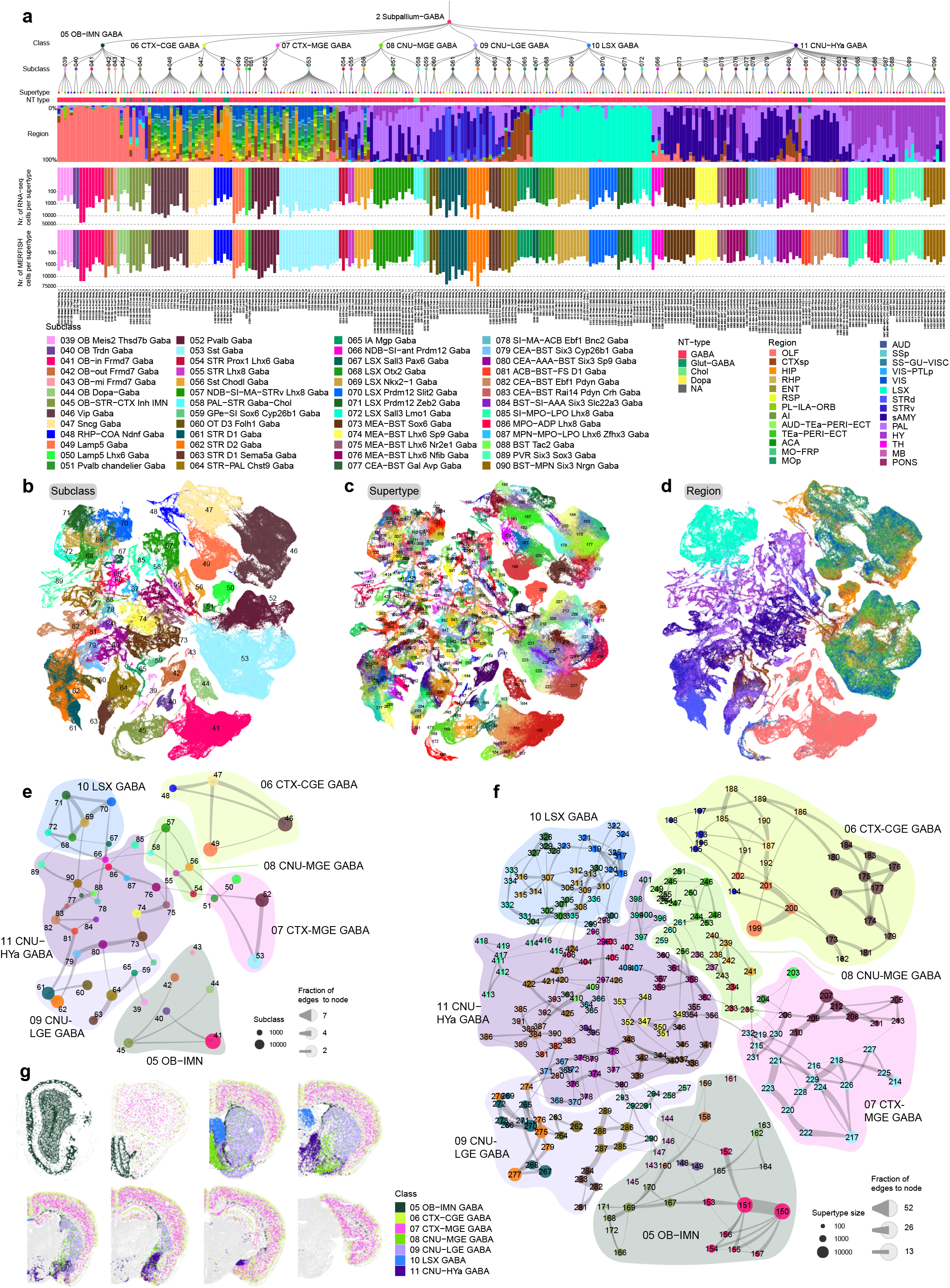
Transcriptomic taxonomy of telencephalic GABAergic neuronal types in the mouse. **(a)** The transcriptomic taxonomy of 285 supertypes organized in a dendrogram (10xv2: n = 271,656 cells; 10xv3 n = 343,761 cells). From top down, the bar plots represent subclass, major neurotransmitter (NT) type, region distribution of profiled cells, number of RNA-seq cells, and number of MERFISH cells per supertype. **(b-d)** UMAP representation of all cell types colored by subclass **(b)**, supertype **(c)**, and dissection region **(d). (e)** Constellation plot of the global relatedness between subclasses. Each subclass is represented by a disk, labeled by the subclass ID, and positioned at the subclass centroid in UMAP coordinates shown in panel b. The size of the disk corresponds to the number of cells within each subclass, and the edge weights correspond to the fraction of shared neighbors (see **Methods**) between subclasses. (**f**) Constellation plot as in panel e but showing relatedness between supertypes. Each supertype is colored by the subclass it belongs to. Bubbles drawn around supertypes outline the major classes. **(g)** Representative MERFISH sections of adult mouse brain across forebrain structures colored by cell class. Each class is labelled by its ID and shown in the same color in the dendrogram and bubbles in the constellation plot.

As the MERFISH data was registered to the Allen Mouse Brain CCFv3, we could determine the GABAergic neuronal cell type composition, at the supertype level, for each of the telencephalic regions (**Extended Data Figure 1a**). This analysis showed that most supertypes span multiple neighboring regions, except for supertypes located in LSX and MOB-AOB which are located in one dominant region. We further used the Gini coefficient and Shannon diversity index to measure the extent of variation in spatial distribution among supertypes (**Extended Data Figure 1a**), and both reveal high inequality (that is, highly localized patterns) in spatial distribution of each neuronal subclass, with the exception of GABAergic neurons in isocortex which are distributed across most cortical regions.

As expected, vast majority of the clusters are purely GABAergic types (**Figure 1a, Supplementary Table 3**). However, we also identified glutamate-GABA co-releasing types expressing *Slc17a8* in 11 clusters across several subclasses (**Figure 1a, Supplementary Table 3**), as well as low-grade expression of both *Chat* and *Slc18a3* in several clusters in the 46 Vip Gaba subclass, though the expression at cluster level did not cross our threshold to label these clusters as cholinergic. We did identify 11 cholinergic neuron clusters in the 58 PAL-STR Gaba-Chol subclass with complex GABA and/or glutamate co-release patterns (see below), and 2 GABA-cholinergic clusters in the 69 LSX Nkx2-1 Gaba subclass (**Figure 1a, Supplementary Table 3**). We also identified 4 GABA-dopamine co-releasing clusters in the 44 OB Dopa-Gaba subclass (**Figure 1a, Supplementary Table 3**).

Furthermore, we have identified 26 neuropeptide genes that are differentially expressed among the GABAergic types of the telencephalon, many of which had been used as markers for various cell types in previous studies. Most neuropeptides show very restricted expression patterns, like *Edn1* which is mostly restricted to the 52 Pvalb Gaba subclass, and some are more broadly expressed, like *Penk* and *Pnoc* which are expressed in various subclasses within each of the GABAergic cell classes **(Extended Data Figure 2)**. Precise association of these neuropeptides with transcriptomic cell types defined here provides a much-needed link to clarify the transcriptomic identities of cell populations labeled by individual marker genes, as well as to explore the function of these neuropeptides in specific cell types.

### GABAergic neuronal types in the olfactory bulb

GABAergic neuronal types in MOB and AOB are thought to be derived from LGE during development and migrate to the olfactory bulb^9^. Furthermore, OB GABAergic neurons continue to be generated in the adult brain through neurogenesis in the subventricular zone (SVZ) and migration into OB via the rostral migratory stream (RMS)^15–17^. In the OB-IMN GABA class, we defined six OB GABAergic subclasses, 39 OB Meis2 Thsd7b Gaba, 40 OB Trdn Gaba, 41 OB-in Frmd7 Gaba, 42 OB-out Frmd7 Gaba, 43 OB-mi Frmd7 Gaba, and 44 OB Dopa-Gaba, and one GABAergic immature neuronal subclass, 45 OB-STR-CTX Inh IMN (**Extended Data Figure 3**).

Subclass 45 OB-STR-CTX Inh IMN contains immature neurons originating from the subventricular zone that migrate to the OB (**Extended Data Figure 3a-c,k, Extended Data Figure 4a,b**). Within this subclass, 7 supertypes have been defined. Supertypes 166, 168, 171, and 172 are the most immature based on gene expression, spatial location, and their pseudo-temporal ordering^14,18^. The other three supertypes seem to be transition types and show similarity to specific mature subclasses (**Extended Data Figure 3a-c**). Supertype 170 OB-STR-CTX Inh IMN_5 forms a transition from immature neurons to subclass 39 OB Meis2 Thsd7b Gaba.

Neurons in this subclass are located mostly in the internal plexiform and mitral (Ipl/Mi), external plexiform (EPl), and glomerular (Gl) layers (**Extended Data Figure 3d,e**). Supertype 143 within the 39 OB Meis2 Thsd7b Gaba subclass are the *Calb1*-positive Blanes cells^19,20^ that populate the Gl (**Extended Data Figure 3e**). Immature neuron supertype 167 OB-STR-CTX Inh IMN_2 forms a transition to the 41 OB-in Frmd7 Gaba subclass. Neurons in this subclass represent the granule cells which populate the granule layer (GrO), IPl/Mi, and supertype 150 contains a subpopulation of neurons that extends into the EPl (**Extended Data Figure 3j)**. Supertype 150 corresponds to a population of glomerular cells that undergoes expansion during periods of olfactory enrichment and contraction during periods of olfactory deprivation (**Extended Data Figure 4a**)^21^. Lastly, supertype 169 OB-STR-CTX Inh IMN_4 forms a transition from immature neurons to subclass 42 OB-out Frmd7 Gaba. Neurons in this subclass exclusively populate the Gl and represent the *Calr*-positive periglomerular cells (PGC) (**Extended Data Figure 3g**).

The 40 OB Trdn Gaba subclass contains the population of *Rprm*-positive granule cells (**Extended Data Figure 3c,f**) that have been identified previously^21,22^. The 43 OB−mi Frmd7 Gaba subclass represents *Pvalb*-positive GABAergic OB neurons described by Batisto-Brito et al.,^20^ which are postnatally generated neurons populating the EPl (**Extended Data Figure 3h**). The glomerular layer of the olfactory bulb signifies the location where sensory input from the olfactory epithelium is first processed within the CNS and contains a variety of interneuron populations which can broadly be divided into three categories; *Calb1*-positive PGC present in subclass 39, *Calr*-positive PGC present in subclass 42, and dopaminergic PGC marked by expression of *Slc6a3* and *Th* present in subclass 44 ^20,21,23–25^ (**Extended Data Figure 3c,e,g,i, Extended Data Figure 4a**).

### GABAergic neuronal types in the cerebral cortex

The cerebral cortex, also known as the pallium, includes four major structures – isocortex, hippocampal formation (HPF), olfactory areas (OLF) and cortical subplate (CTXsp). The GABAergic neurons in all these brain structures belong to two classes, CTX-CGE GABA (**Extended Data Figure 5**) and CTX-MGE GABA (**Figure 2**), named after their spatial location and dominant developmental origins^5,12^. The current classification is largely consistent with our previously defined transcriptomic taxonomy of the mouse isocortex and HPF^26^ as well as the MET-types in our Patch-seq study in the mouse visual cortex^27^ (**Extended Data Figure 6**), and further extends into the OLF and CTXsp areas.

**Figure 2.**
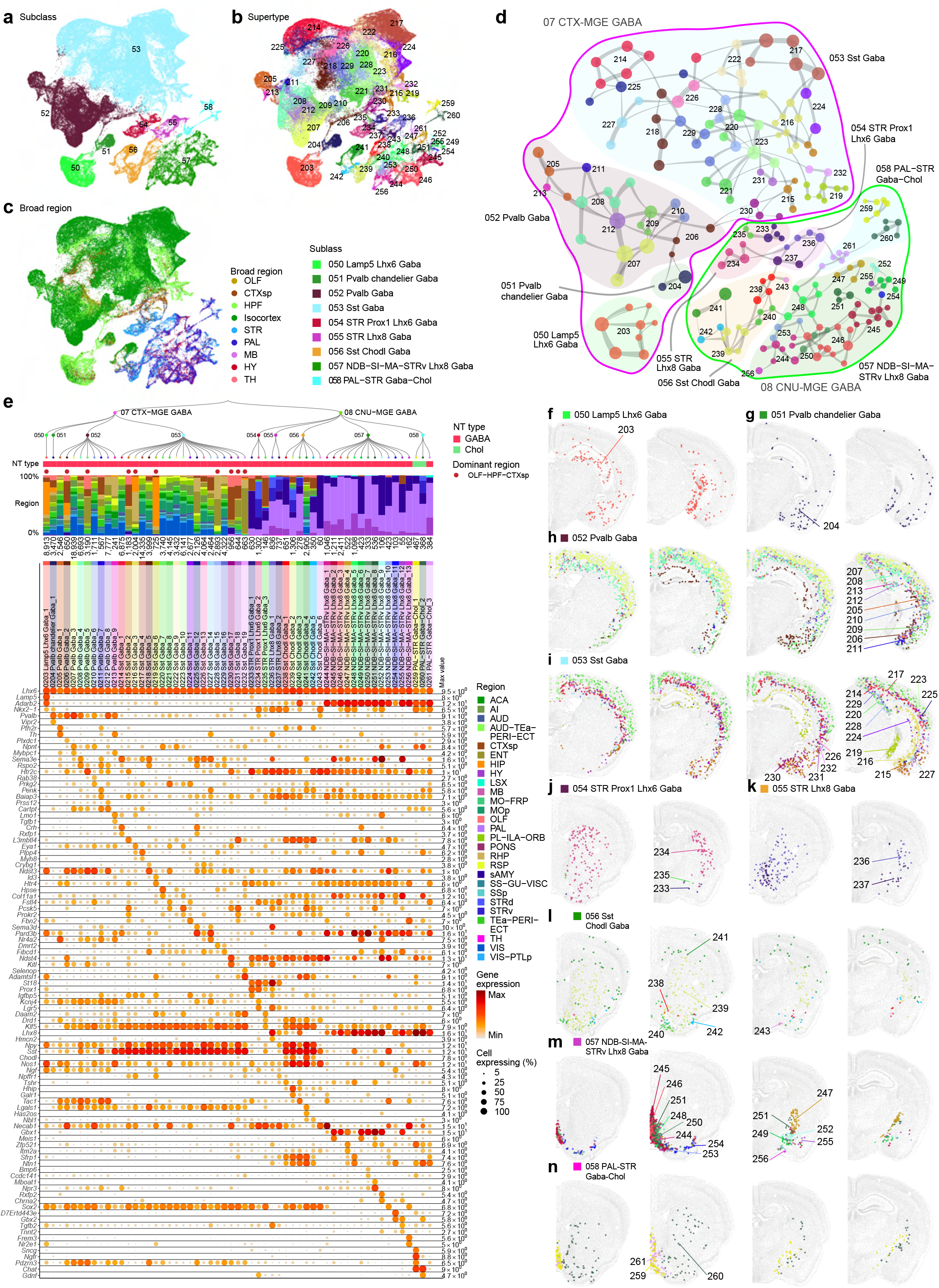
MGE-derived GABAergic cell types in the cerebral cortex and cerebral nuclei. **(a-c)** UMAP representation of all MGE clusters colored by subclass (a), supertype (b), or broad brain region (c). **(d)** Constellation plot of MGE clusters using UMAP coordinates shown in b. Nodes are colored by supertype and grouped in bubbles by subclass. Lines around the bubbles denote the class the nodes belong to. **(e)** Dendrogram of MGE supertypes followed by bar graphs showing major neurotransmitter type, region distribution of profiled cells, dominant region, and number of cells within supertype, followed by dot plot showing marker gene expression in each supertype from the 10xv3 dataset. The dominant region was assigned if more than 70% of cells are from the OLF-HPF-CTXsp regions. For the gene expression dot plot, dot size and color indicate proportion of expressing cells and average expression level in each supertype, respectively. **(f-n)** Representative MERFISH sections showing the location of supertypes in MGE subclasses 50 Lamp5 Lhx6 Gaba (f), 51 Pvalb chandelier Gaba (g), 52 Pvalb Gaba (h), 53 Sst Gaba (i), 54 STR Prox1 Lhx6 Gaba (j), 55 STRv Lhx8 Gaba (k), 56 Sst Chodl Gaba (l), 57 NDB−SI−MA−STRv Lhx8 Gaba (m), 58 PAL−STR Gaba−Chol (n). Cells are colored and labelled by supertype.

In the CTX-CGE GABA class, we defined four subclasses, including three previously defined ones – 46 Vip Gaba, 47 Sncg Gaba and 49 Lamp5 Gaba^26,28^, and one newly defined here, 48 RHP-COA Ndnf Gaba (**Extended Data Figure 5a-g**). The Vip, Sncg and Lamp5 subclasses correspond to the well-known bipolar, multipolar, neurogliaform and other types of GABAergic interneurons widely distributed in all areas of isocortex and HPF. Here we find that they are also present in OLF and CTXsp areas (**Extended Data Figure 5c-g**). Consistent with previous findings, most of their clusters and supertypes are shared among all pallial areas, whereas 8 supertypes are predominant in OLF/CTXsp/HPF areas (marked by red dots, **Extended Data Figure 5c**). We also identified a set of Vip Gaba clusters that are largely specific to hippocampus (HIP), including clusters 649, 650, 651, 654, 655, and 659, included in supertypes 179, 181, and 182 respectively (**Extended Data Figure 5b-d**).

The newly defined RHP-COA Ndnf Gaba subclass expresses marker genes *Ndnf* and *Ntng1*, is predominantly present in HPF, OLF and CTXsp and exhibits high regional specificity. In this subclass, supertypes 195 and 198 are mainly found in ENT and RHP respectively, supertype 194 is located in COA, and remaining supertypes are mainly present in HIP (**Extended Data Figure 5c,f**). The RHP-COA Ndnf Gaba subclass also includes the previously described *Meis2*-positive population of GABAergic neurons in supertype 198 (**Extended Data Figure 5a-c**)^26,28^. These *Meis2*-positive GABAergic neurons reside in the white matter and originate from the embryonic pallial-subpallial boundary^29,30^. The cells in 198 supertype are the only cells in the cortical GABAergic classes that express *Meis2* (**Extended Data Figure 5c**). However, the entire subclass lacks expression of *Prox1* which is found in all other CGE-derived GABAergic neurons and shares expression of *Rspo3* and *Ntng1*.

In the CTX-MGE GABA class, we defined four subclasses, including three previously defined ones^26,28^, 53 Sst Gaba, 52 Pvalb Gaba and 51 Pvalb chandelier Gaba, and the fourth one, 50 Lamp5 Lhx6 Gaba, which was previously considered as part of the CGE Lamp5 subclass but is now classified into the MGE class based on its expression of the MGE transcription factor, *Lhx6* (**Figure 2a-d**). The Sst, Pvalb and Pvalb chandelier subclasses correspond to the well-known somatostatin, parvalbumin, and chandelier types of GABAergic interneurons widely distributed in all areas of isocortex and HPF. Here we find that they are also present in OLF and CTXsp areas (**Figure 2d-i**). There are two Pvalb chandelier clusters, one widely distributed in all pallial areas whereas the other is mainly found in HIP. In the Pvalb GABA subclass, individual clusters display more variable regional preference, with 4 supertypes mainly in isocortex, and 2 supertypes mainly in OLF/CTXsp/HPF out of 9 supertypes in total (**Figure 2e**). The Sst GABA subclass is highly diverse, with a total of 19 supertypes and 71 clusters, and variable regional preference. Seven supertypes are predominant in OLF/CTXsp/HPF areas (marked by red dots, **Figure 2e**) whereas the other supertypes are more broadly present in isocortex. Several Pvalb and Sst supertypes are largely specific to HPF, including 209, 216, 219, 228, and 232 (**Figure 2e,h,i**). The Lamp5 Lhx6 subclass is predominantly present in HIP, with a small fraction of cells found in other pallial areas (**Figure 2e,f**).

Overall, the organization of GABAergic neurons in the cerebral cortex (pallium) exhibits a clear segregation of cell types (in both CTX-CGE and CTX-MGE classes) between isocortex (also known as the neocortex) and the other structures that are considered evolutionarily more ancient, including HPF, OLF, and CTXsp. We observe a gradual transition between these two parts, with certain isocortical areas as intermediates, e.g., retrosplenial cortex (RSP). For example, supertype 177 Vip Gaba_5 is mostly located to isocortex and via RSP shows sparse labeling in HIP (**Extended Data Figure 5d**).

### MGE-derived GABAergic neuronal types in the cerebral nuclei

GABAergic neurons originating from the MGE show wide-spread distribution. These MGE-derived neurons are divided into two main classes; where the CTX-MGE class located mostly in pallial areas as described above, the CNU-MGE located mainly in the cerebral nuclei. These two classes share the expression of *Lhx6*, a developmental pan-MGE marker. There is a striking difference in complexity between the two MGE classes. The CNU MGE-derived GABAergic neurons form clusters that are typically smaller and more heterogeneous than the CTX MGE-derived GABAergic neurons (**Figure 2a-d**).

The cerebral nuclei (CNU) are composed of two major structures, striatum (STR) and pallidum (PAL). Its neuronal population is largely GABAergic. The GABAergic neurons in CNU come from four classes (**Figure 1**). The CNU-MGE GABA and CNU-LGE GABA classes contain striatal and pallidal GABAergic neurons derived from MGE and LGE respectively. The LSX GABA class contains lateral septum GABAergic neurons derived from the embryonic septum^31^. The CNU-HYa GABA class contains GABAergic neurons mainly located in sAMY, PALc and anterior hypothalamus (**Extended Data Figure 1a**); these neurons may also be developmentally originated from LGE, MGE, as well as the embryonic preoptic area (POA).

In the CNU-MGE class, we defined 5 subclasses which are 54 STR Prox1 Lhx6 Gaba, 55 STR Lhx8 Gaba, 56 Sst Chodl Gaba, 57 NDB-SI-MA-STRv Lhx8 Gaba, and 58 PAL-STR Gaba-Chol (**Figure 2a-e**). These subclasses are mainly located in dorsal and ventral striatum (STRd and STRv), and dorsal, ventral and medial pallidum (PALd, PALv and PALm). Each subclass is not specific to a single region but contains neurons from multiple regions (**Figure 2j-n**). Although local interneurons make up less than 10% of the striatal neurons, these cells represent a diverse population (**Extanded Data Figure 4d**)^32,33^. The CNU-MGE types located within the STRd and STRv are likely striatal GABAergic interneurons, including supertypes 234 STR Prox1 Lhx6 Gaba_2 (*Pvalb*-positive) and 236 STRv Lhx8 Gaba_1 (*Pvalb*-negative). Interestingly, the 56 Sst Chodl subclass is a unique subclass that spans both pallium and subpallium structures, containing neurons in isocortex (supertype 241, the Sst Chodl cells^26,28^), CTXsp (supertype 242), STRd/STRv, and PALv/PALc (**Figure 2d,e,l, Extended Data Figure 1a**).

### LGE-derived GABAergic neuronal types in the cerebral nuclei

The seven LGE-derived subclasses in the cerebral nuclei are 59 GPe Sox6 Cyp26b1 Gaba, 60 OT D3 Folh1 Gaba, 61 STR D1 Gaba, 62 STR D2 Gaba, 63 STR D1 Sema5a Gaba, 64 STR-PAL Chst9 Gaba, and 65 IA Mgp Gaba (**Figures 3**). These subclasses resemble the well-known D1 and D2 type medium spiny neurons (MSN; also known as spiny projection neurons, SPN) in the striatum, with the 61 STR D1 Gaba and 62 STR D2 Gaba subclasses being the prototype striatal D1 and D2 cell types and the other subclasses being newly defined homologous cell types. Among them, subclasses 60, 63 and 64 also express dopamine receptor gene *Drd1*, and subclass 60 additionally expresses *Drd3* strongly (**Extended Data Figure 7a**).

**Figure 3.**
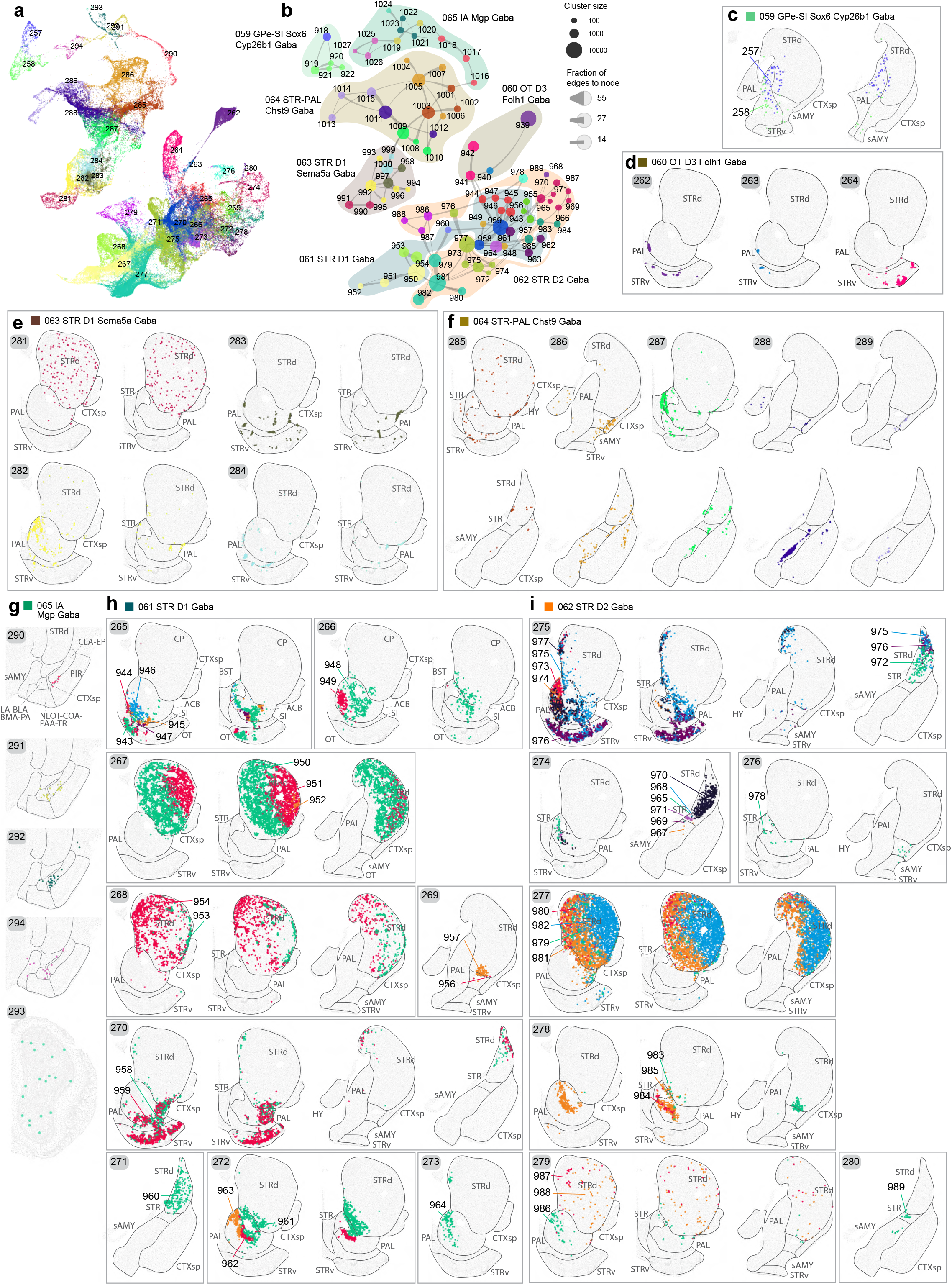
LGE-derived GABAergic cell types of the cerebral nuclei. **(a)** UMAP representation of all LGE clusters colored by supertype. **(b)** Constellation plot of LGE clusters using UMAP coordinates shown in a. Nodes are colored by supertype and grouped in bubbles by subclass. **(c-g)** Representative MERFISH sections showing the location of LGE subclasses, 58 GPe Sox6 Cyp26b1 Gaba (c), 60 OT D3 Folh1 Gaba (d), 63 STR D1 Sema5a Gaba (e), 64 STR−PAL Chst9 Gaba (f), 65 IA Mgp Gaba (g). Cells are colored and labelled by supertype. **(h-i)** Representative MERFISH sections showing the location of CNU-LGE subclasses, 61 STR D1 Gaba (h) and 62 STR D1 Gaba (i). Each box contains one supertype and cells are labelled and colored by cluster to highlight the diversity.

Subclass 60, with its strong expression of Drd3 along with Drd1, is specifically localized in the islands of Calleja in the OT (**Figure 3d, Extended Data Figure 4c,d**). Subclasses 63, 64 and 65 collectively form a novel group of GABAergic neuronal types that are like MSNs based on their gene expression profile but also distinct from them based on their spatial location (**Figure 3e-g**). The spatial distribution patterns of many supertypes and clusters in these subclasses are remarkably unique, widely scattered along the borders between different striatal and pallidal areas, forming multiple patches or streaks (**Figure 3e-g**). Subclass 63 STR D1 Sema5a Gaba contains hybrid MSN D1 neurons that have been described to co-express *Drd1a* and a shortened variant of the D2 receptor (**Figure 3e**, **Extended Data Figure 4c**)^34,35^. This group of MSN resembles so-called exopatch MSN, which are located in the striatal matrix but physiologically resemble MSN of patches (**Extended Data Figure 4d**)^33,36^. Based on their spatial distribution patterns, we assigned the locations of subclasses 64 STR-PAL Chst9 Gaba and 65 IA Mgp Gaba to interstitial nucleus of the posterior limb of the anterior commissure (IPAC, described in Paxinos’s atlas^37^) or intercalated amygdalar nucleus (IA). Subclass 65 is divided into five different supertypes which all express *Dcx*, a migratory neuroblast marker, and markers *Ncam1* (PSA-NCAM), *Foxp2* and *Meis2*, indicating that these are immature neurons. In addition to

*Foxp2*, these types express *Tshz1*, which are known markers for the intercalated cells originating from the *Sp8*-positive dLGE progenitor population^38–40^. Interestingly, we observed a relatedness between subclass 65 IA Mgp Gaba and subclass 39 OB Meis2 Thsd7b Gaba (**Figure 1e,f**), suggesting that they may have shared developmental origins. Within subclass 65, supertype 293 stands out in that it is spatially located in the olfactory bulb with sparse labeling in IA (**Figure 3g**), highlighting the similarity between the intercalated cells and the immature neurons migrating towards the olfactory bulb.

We defined 9 supertypes in 61 STR D1 Gaba subclass and 7 supertypes in 62 STR D2 Gaba subclass. The supertypes and clusters in both subclasses exhibit highly diverse spatial distribution patterns in STRd and STRv (**Figure 3h,i**). Some occupy the entire space, while others are specifically located in medial-lateral, anterior-posterior, and dorsal-ventral subdomains. Supertype 268 STR D1 Gaba_4 within the 61 STR D1 Gaba subclass is mostly located in striosomes, while supertype 267 STR D1 Gaba_3 is mostly located in the striatal matrix (**Figure 3h**). Similarly, within the 62 STR D2 Gaba subclass, supertype 279 STR D2 Gaba_6 is mostly present in the striosomes, while supertype 277 STR D2 Gaba_4 represents types in the striatal matrix (**Figure 3i**).

We found complex subregional enrichment of cell types in the nucleus accumbens (ACB). Supertypes 265, 266, 270, 272, and 273 within the 61 STR D1 Gaba subclass and supertypes 275 and 278 within the 62 STR D2 Gaba subclass are mostly restricted to ACB and OT (**Figure 3h,i**). The ACB can be divided into core and shell subregions^41,42^. Supertypes 272 STR D1 Gaba_8, 273 STR D1 Gaba_9, and 278 STR D2 Gaba_5 are predominantly located in the core region. Supertype 266 STR D1 Gaba_2 consists of two highly related cell types of which cluster 948 is in the core subdomain and cluster 949 is located in the shell subdomain (**Figure 3h**). Similarly, supertype 275 STR D2 contains various clusters of which 973 and 974 are predominantly located in the core subdomain and the other types are present in the shell subdomain or OT (**Figure 3i**). Based on the cell types and their locations the core and shell regions of ACB can be further divided on a mediolateral and anteroposterior axis. For example, supertype 272 STR D1 Gaba_8 consists of three different clusters, with cluster 963 located in the medial-anterior subdomain of the ACB core while cluster 961 in a more lateral-posterior position (**Figure 3h**). The same can be seen for ACB core D2 neurons in supertype 278 STR D2 Gaba_5, with cluster 985 in a medial-anterior position and cluster 983 in a more lateral-posterior position. Similar distribution of cell types along the different anatomical axes can be observed for the types in the shell subdomain of ACB. Within the STR D1 Gaba subclass, the shell cell types are present in supertypes 265 STR D1 Gaba_1 and 270 STR D1 Gaba_7 and occupy medial and lateral subdomains of shell, respectively. Compared to the lateral types (supertype 270), the medial types (supertype 265) show higher diversity and distinct distribution along the anteroposterior axis (**Figure 3h**). As observed previously, though the cell types are preferentially located in either core or shell domains of ACB, the cell types occupy overlapping regions^33^.

### GABAergic neuronal types in the lateral septum of the cerebral nuclei

The LSX GABA class is located specifically in lateral septal complex (LSX) and is highly distinct from other CNU GABAergic neuronal types. It can be divided into six subclasses, 67 LSX Sall3 Pax6 Gaba, 68 LSX Otx2 Gaba, 69 LSX Nkx2-1 Gaba, 70 LSX Prdm12 Slit2 Gaba, 71 LSX Prdm12 Zeb2 Gaba, and 72 LSX Sall3 Lmo1 Gaba, but the relationships among these subclasses are complex and intertwined (**Extended Data Figure 8a-c**). The LSX subclasses exhibit partially overlapping spatial distribution patterns within the lateral septum (**Extended Data Figure 8d-j**). For example, the 67 LSX Sall3 Pax6 Gaba and 68 LSX Otx2 Gaba subclasses are both present in rostral (LSr) and ventral (LSv) regions of LSX, with the 68 LSX Otx2 extending more into dorsal LSX (**Extended Data Figure 8d-f**). Another example is subclasses 69 LSX Nkx2-1 Gaba and 70 LSX Prdm12 Slit2 Gaba that have shared presence in LSr and LSv, with 69 extending more into anterior LSX and 70 extending into posterior LSX (**Extended Data Figure 8d,g,h**).

### GABAergic neuronal types in the striatum-like amygdalar nuclei

The CNU-HYa GABA class is highly complex, with 19 subclasses that are predominantly localized in CNU but also extend into the preoptic area (POA) of the anterior hypothalamus. The 19 subclasses are 66 NDB-SI-ant Prdm12 Gaba, 73 MEA-BST Sox6 Gaba, 74 MEA-BST Lhx6 Sp9 Gaba, 75 MEA-BST Lhx6 Nr2e1 Gaba, 76 MEA-BST Lhx6 Nfib Gaba, 77 CEA-BST Gal Avp Gaba, 78 SI-MA-ACB Ebf1 Bnc2 Gaba, 79 CEA-BST Six3 Cyp26b1 Gaba, 80 CEA-AAA-BST Six3 Sp9 Gaba, 81 ACB-BST-FS D1 Gaba, 82 CEA-BST Ebf1 Pdyn Gaba, 83 CEA-BST Rai14 Pdyn Crh Gaba, 84 BST-SI-AAA Six3 Slc22a3 Gaba, 85 SI-MPO-LPO Lhx8 Gaba, 86 MPO-ADP Lhx8 Gaba, 87 MPN-MPO-LPO Lhx6 Zfhx3 Gaba, 88 BST Tac2 Gaba, 89 PVR Six3 Sox3 Gaba, and 90 BST-MPN Six3 Nrgn Gaba (**Extended Data Figure 9**). Neurons in this class are located in specific sAMY and PAL areas, including central amygdalar nucleus (CEA), anterior amygdalar area (AAA), bed nuclei of the stria terminalis (BST), substantia innominata (SI), and magnocellular nucleus (MA) (**Figure 4a**, **Extended Data Figure 9d**). Each subclass within the CNU-HYa GABA class is not specific to a single region but contains neurons from multiple regions. For example, subclass 80 contains neurons from CEA, AAA, and BST (**Figure 4a,c**, **Extended Data Figure 9d**). Conversely, each region contains multiple subclasses. For example, subclasses 74,79, 80, and 82 are co-localized in CEA, AAA, and BST. Subclasses 78 and 84 are co-localized in SI, NDB, and MA.

**Figure 4.**
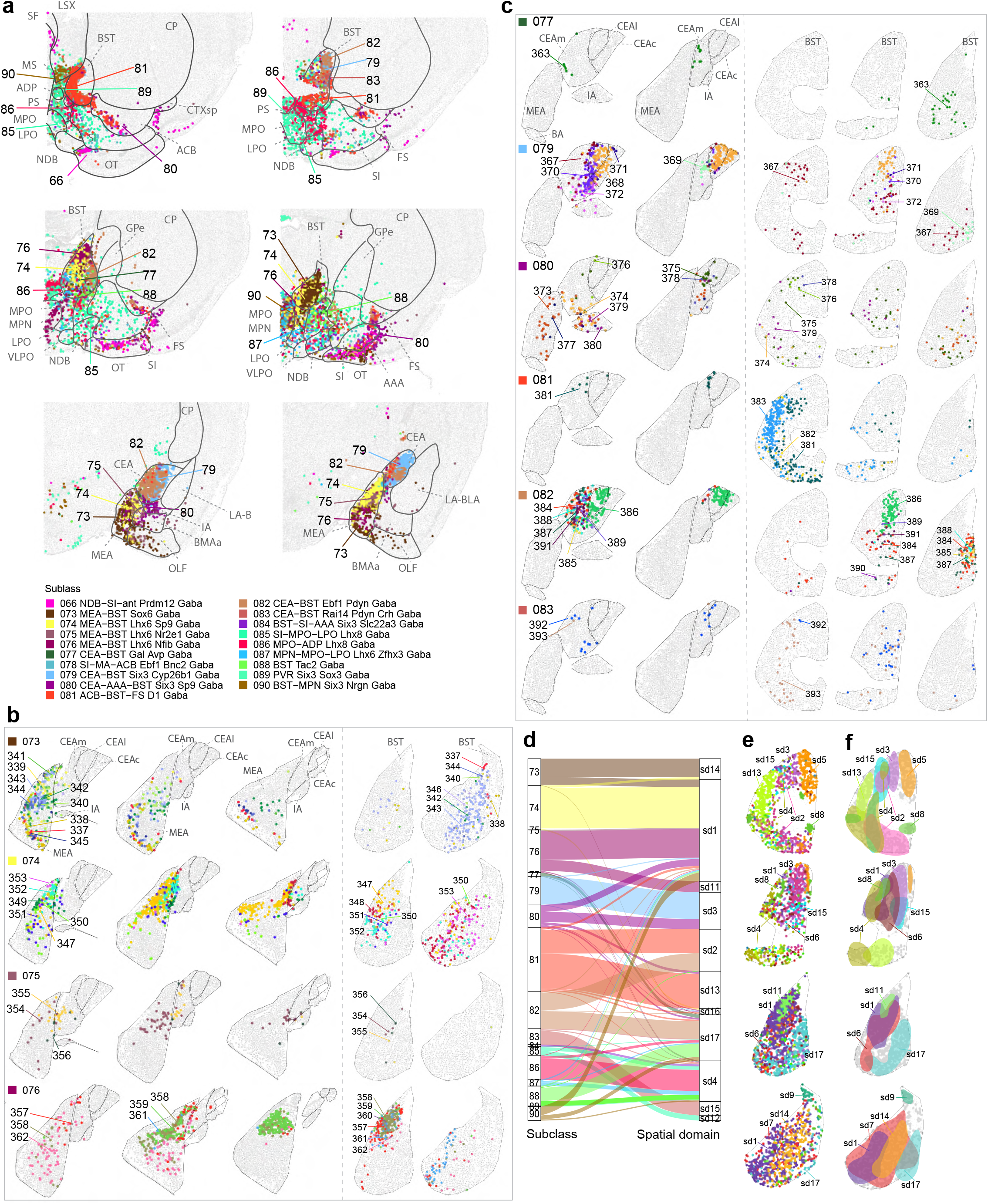
Organization of GABAergic neuronal types across striatum-like amygdalar nuclei and bed nuclei of the stria terminalis. **(a)** Representative MERFISH sections showing the location of subclasses belonging to the CNU-HYa neuronal class. Cells are colored by the subclass they belong to and labelled by its ID. **(b-c)** Representative MERFISH sections showing the locations of cells belonging to the MEA-BST subclasses (b) and CEA-BST subclasses (c). Each row shows one subclass, and cells are colored and labelled by supertype identity. For each subclass the location in both MEA and BST (b) or CEA and BST (c) are shown, indicating the existence of the same cell types in these locations. **(d-f)** Spatial domain clustering within BST neurons using Banksy. The alluvial plot (d) shows the relation between subclasses present in BST and the spatial domains. Representative MERFISH images show the location of the spatial domains, with cells colored by spatial domain identity (e) and the area covered by each domain (f).

BST serves as a hub for processing limbic information and monitoring emotional valence and the center of a vast connectivity network. Mood and arousal are processed via connections to sAMY, dorsal raphe, and the ventral tegmental area (VTA). Via connections to the hypothalamus, BST monitors feeding and drinking signals coming from brainstem. Via connections to lateral septum and MEA, BST coordinates reproductive and related social behaviors. The diverse connectivity patterns in which the BST participates appear to be associated with specific cell types. BST, also referred to as part of the extended amygdala, is a major output region for neurons in both CEA and MEA^43–45^. Our data shows a high transcriptomic similarity of cell types in CEA and BST, and of cell types in MEA and BST, while these two groups are easily distinguishable by distinct marker genes (**Extended Data Figure 9d**).

Types within subclasses 73, 74, 75, 76, and 88 are mostly located in MEA and/or BST (**Figure 1a,e**, **Figure 4b, Extended Data Figure 9d**), and possibly share a similar developmental origin based on preserved transcription factor expression (see below). The MEA-BST types also show similarity to subclasses 85, 86, 87, 89 and 90 which are mostly located in POA and are known to share functions and circuits^46–49^ (**Figure 1e, Extended Data Figure 9d**). MEA plays a central role in regulating social behavior in rodents. It is the location where signals from the OB and vomeronasal system converge, placing the MEA in a position critical for processing pheromonal signals that regulate social behavior^50,51^. For example, subclasses 74, 75, 76 express the MEApd marker *Lhx6*^52,53^ and neurons in this region are involved in reproductive behaviours^50,53,54^. Supertypes 349 and 351 within subclass 74 MEA-BST Lhx6 Sp9 Gaba express *Crhr2* and *Ucn3* (**Extended Data Figure 2**). Cell types expressing these genes have been identified as part of the behavioral stress response system^55^. Supertypes 357 to 359 within subclass 76 MEA-BST Lhx6 Nfib Gaba contain the BNSTpr^Tac^^1/Esr^^1^ cell type that is essential for male social interactions (**Extended data Figure 10a**)^49^.

Subclasses 77 to 84 are mostly located in CEA, BST, AAA and SI (**Extended Data Figure 9d**), are similar to striatal CNU-LGE subclasses (**Figure 1e**, **Figure 4a,c**), and possibly share a similar developmental origin based on preserved transcription factor expression (see below). CEA is a striatal-like GABAergic structure that contains both GABAergic interneurons and GABAergic long-range projection neurons and projects to the hypothalamus and brainstem to initiate fear responses^56^. Most supertypes in subclass 79 CEA-AAA-BST Six3 Cyp26b1 Gaba are located in the lateral and capsular part of CEA (CEAl and CEAc), express known MSN markers *Penk*, *Pax6*, *Gpr88*, and *Ppp1r1b*^34,35,52^, and are related to subclass 62 STR D2 Gaba (**Figure 1e,f**, **Figure 4c, Extended Data Figure 9d**). Supertype 371 CEA-AAA-BST Six3 Cyp26b1 Gaba_5, located mostly in CEAc, is closely related to supertypes 274 STR D2 Gaba_1 and 280 STR D2 Gaba71 (**Figure 1e,f**, **Figure 4c**). Also based on their spatial location these supertypes are in close proximity to each other (**Figure 3i**). Supertype 368 CEA-AAA-BST Six3 Cyp26b1 Gaba_2 corresponds to the previously described CeA Prkcd-Ezr type that is highly responsive to cued fear conditioning (**Extended Data Figure 10b**)^52^. Subclasses 77 CEA-BST Gal Avp Gaba, 82 CEA-BST Ebf1 Pdyn Gaba and 83 CEA-BST Rai14 Pdyn Crh Gaba are mainly located in the medial part of CEA (CEAm, **Figure 4c**).

As the MERFISH data was registered to the Allen Mouse Brain CCFv3, we could select all neurons within BST from the MERFISH dataset and apply spatial clustering to identify subdomains. Cells were clustered on both their gene expression pattern based on the 500 gene panel and their spatial location within the neighborhood (see **Methods**). Using this approach, we identified unique domains of which several align with previously described anatomical subdivisions of BST, and these align with the MEA-BST and CEA-BST cell subclasses (**Figure 4d**). Several subdivisions of BST have been proposed; a mediolateral division based on cytoarchitecture and input connections from amygdala, an anteroposterior division based on developmental origin, and a dorsoventral division based on divergent patterns of monoaminergic innervation^43,57–61^. MEA projects mostly to posteromedial BST regions^43^ which is an area where the MEA-BST subclasses (73-76) are enriched. CEA, on the other hand, mostly projects to anterolateral regions in BST^62^ which is where we see enrichment of CEA-BST subclasses (77, 79-83) (**Figure 4e-f**).

### Long-range GABAergic projection neurons

Most GABAergic neurons in the cerebral cortex are local interneurons, except for a small set of cell types that have long-range projections (LRP)^63–65^. The LRP types in the hippocampus (HIP) are among the best described cortical LRP neurons. Based on expression of genes, including *Chrna4*, *Pcp4*, *Nos1* and *Htr3a*, supertype 179 Vip Gaba_7, located in HIP, is marked as a LRP population^66^. The 215 Sst Gaba_2 and 216 Sst Gaba_3 supertypes contain putative LRP neurons expressing *Sst*, *Npy,* and *Nos1*, and located in OLF/CTXsp regions or HPF regions, respectively (**Figure 2i**).

The previously identified Sst Chodl long-range projecting neurons in cortex^27,28,67,68^ are closely related to the local Sst interneurons in CNU, as they all belong to a single subclass, 56 Sst Chodl Gaba, based on their gene expression profiles (**Figure 2a,d,e,l**). Supertype 241 Sst Chodl Gaba_4 contains the cortical Sst Chodl neurons whereas the other 5 supertypes in this class are mostly located in CNU.

The 58 PAL-STR Gaba-Chol subclass contains basal forebrain cholinergic neurons (**Extended Data Figure 11**). This subclass is divided into 3 supertypes, with supertypes 259 and 260 containing cholinergic neurons. The third one, supertype 261, contains closely related GABAergic neurons that are mainly located in medial septum (MS) and diagonal band nucleus (NDB). The cholinergic neurons have cluster-level specificity in their spatial localizations. Those in supertype 259 are located in MS and NDB (clusters 923-925), substantia innominata (SI) and NDB (clusters 926 and 928), or GPe and SI (cluster 927) (**Extended Data Figure 11b**), and thus are assigned as the cholinergic projection neurons in the basal forebrain. Those in supertype 260 are located in STR (including CP, ACB, OT, etc.) (**Extended Data Figure 11c**), and thus are assigned as the striatal cholinergic interneurons. All cholinergic projection neuron clusters in supertype 259 co-release GABA, some of which also co-release glutamate. On the other hand, we were surprised to find that none of the striatal cholinergic interneuron clusters in supertype 260 co-release GABA, instead, some of these clusters even co-express glutamate (via transporter gene *Slc17a8*) (**Extended Data Figure 11e, Supplementary Table 3**). Other cholinergic neurons in CNU include clusters 1081 and 1084 within the 69 LSX Nkx2-1 Gaba subclass and 307 LSX Nkx2-1 Gaba_2 supertype, which are both GABAergic and cholinergic (**Extended Data Figure 11a,d,e**).

### Gene expression gradients within telencephalic brain structures

#### MGE cortical gradient

During adulthood, CGE-derived GABAergic neurons in the 46 Vip Gaba and 47 Sncg Gaba subclasses are distributed evenly through either deep or superficial cortical layers and do not show much of a spatial gene expression gradient (**Extended Data Figures 1b, 5c-e**). In contrast, MGE-derived GABAergic neurons adopt their laminar distribution in an ‘‘inside-out’’ manner that correlates with their birthdate, later-born neurons migrate radially past earlier-born neurons to populate more superficial layers. *Sst*-positive GABAergic neurons are among the first types to diversify and mature and are more abundant in infragranular than in supragranular layers of the isocortex, while *Pvalb*-positive GABAergic neurons can be found throughout all layers except Layer 1 (**Figure 2h-i, Extended Data Figure 12a-c**). We performed Independent Component Analysis (ICA) on the Pvalb GABA and Sst GABA classes independently, projected the scRNA-seq result onto the MERFISH data and extracted the top loading genes for the components with the strongest spatial gradient (see **Methods**). We identified 20 genes that drive a spatial gradient along the cortical depth in the Pvalb GABA subclass and 45 genes that drive a spatial gradient in the Sst GABA subclass (**Extended Data Figure 12d-g**). Among these genes, there only five genes that are shared between the subclasses (*Gm32647*, *Il1rapl2*, *Calb1*, *Nkain3*, and *Parm1*) indicating that the observed gradient is unique for each subclass. Among the genes driving the gene expression gradient there are only 4 genes that are subclass or supertype markers (*Rbp4*, *Pdyn*, *Nr2f2*, and *Plpp4*), suggesting that the spatial gene expression gradient is not only driven by supertype diversity within the subclass.

#### Homology and gradients in MSN populations

There exists a strong similarity between STR D1 and STR D2 GABAergic neurons. The constellation plot shows most similar pairs of STR D1 and STR D2 neurons (**Figure 5a**, see **Methods**). We selected five cluster pairs along the UMAP distribution and examined their spatial location using the MERFISH data (**Figure 5b-f**). These closely related STR D1 and STR D2 clusters share the same spatial locations. Next, we performed Independent Component Analysis (ICA) on the STR D1 and STR D2 subclasses independently, projected the scRNA-seq result onto the MERFISH data, and extracted the top common genes among STR D1 and D2 clusters driving the spatial gradient (see **Methods, Figure 5g,h**). We identified two gene modules that represent the dorsolateral-to-ventromedial ends of the gradient. For each cell we calculated the gene module score for each of the two modules and show the scores on both the UMAP based on single cell RNA-seq and representative MERFISH sections (**Extended Data Figure 7b-i**). The results show that a similar spatial gradient drives the dorsolateral-to-ventromedial differences seen in both STR D1 and D2 types. The ventromedial located STR D1 and D2 types are marked by expression of genes coding for various neuroactive receptors, like *Grm8*, *Htr2a*, and *Cnr* (**Figure 5g,h**). The dorsolateral located STR D1 and D2 types on the other hand are marked by expression of genes coding for proteins involved in cGMP-PKG signaling including *Csgalnact1*, *Prkg1*, and *Slc8a1*.

**Figure 5.**
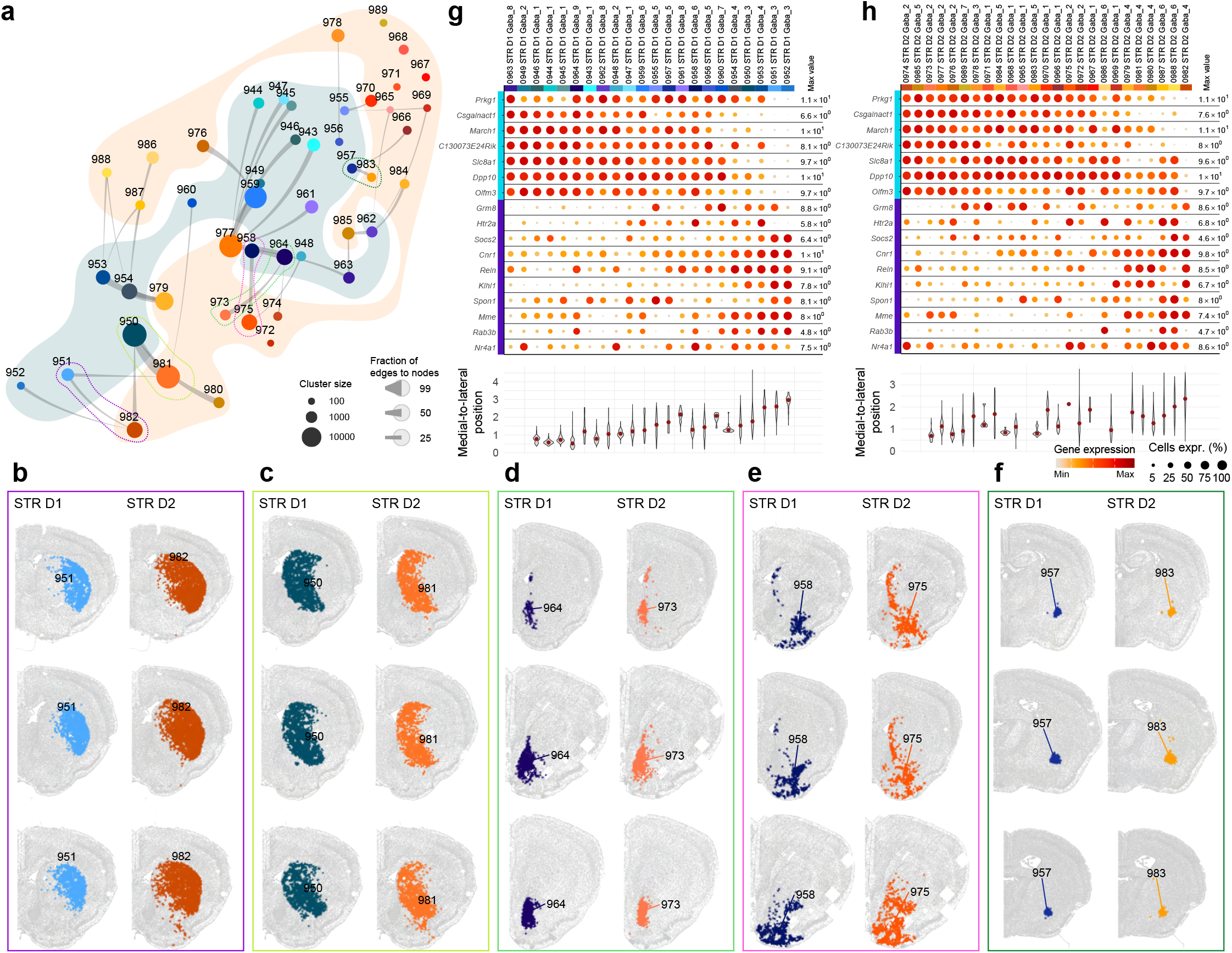
Gene signatures defining shared gradients in D1 and D2 medium spiny neurons. **(a)** Constellation plot of clusters showing the pairs of most similar clusters between 61 STR D1 Gaba and 62 STR D2 Gaba subclasses. Clusters are represented by a disk colored by cluster, labeled by cluster ID, and disks with a colored border are highlighted exemplars in panels b to f. (**b-f**) Representative MERFISH sections showing five examples of STR D1 and STR D2 pairs from panel a, and their spatial distribution patterns. Sections are colored by cluster identity. (**g-h**) Gene expression dot plots of two major gene modules driving the spatial gradient among STR D1 (g) and STR D2 (h) clusters. Dot size and color indicate proportion of expressing cells and average expression level in each cluster, respectively. Underneath the dot plot a violin plot of the medial-lateral (x) coordinate for each MERFISH cell per cluster is shown.

#### LSX gradients

LSX is a structure of the basal forebrain that integrates inputs from many cortical and subcortical regions and transmitting the appropriate signals to downstream hypothalamic and midbrain nuclei. As such it plays a central role in the regulation of social behaviors like anxiety and aggression. Most neurons in LSX are GABAergic and contain receptors for a variety of neuromodulators and neuropeptides (**Extended Data Figure 2**). The relationships among subclasses in LSX are complex, show overlapping spatial distribution patterns, and transcriptomic subclasses or supertypes are not necessarily restricted to a single segment of LSX. By imputing scRNA-seq data into the MERFISH brain space, we examined the 3D spatial gradients in LSX more systematically (**Extended Data Figure 13**). We performed ICA on the LSX GABA class and projected the scRNA-seq result onto the MERFISH data (see **Methods**). For the top five spatial gradients identified, we extracted the genes driving the gradient (gene modules) and calculated a gene score for each cell based on both the positive and negative genes in each of the five modules (**Extended Data Figure 13**). Among the genes in the gene modules highlighted, we see two subclass markers, *Zeb2*, and *Six3*, and just one supertype marker, *Foxp2*. Indicating that most genes driving the spatial gradients are crossing subclass and supertype boundaries (**Extended Data Figure 13a**). The strongest spatial gradients in LSX represent dorsoventral or mediolateral gradients but no strong anteroposterior gradient (**Extended Data Figure 13c**). Earlier reports already indicated that LSX input domains do not align well with the molecular organization within LSX but the molecular organization does align with the long range projections^69,70^. For example, gene module 3 contains *Foxp2* and *Ndst4*, genes that are expressed in a subdomain of LSX and neurons in this area have been shown to project to MPO and LPOA regions in the hypothalamus^70,71^. Our data shows that cell types are organized along multidimensional gradients that might align with both input and output domains in LSX.

### Persistent developmental signatures

The telencephalic GABAergic neurons arise mostly from the five principal progenitor domains of the subpallium and from there migrate to populate various regions of the telencephalon. We identified a comprehensive set of transcription factor (TF) marker genes defining the adult neuronal types at class and subclass levels (**Figure 6a**). To verify that the expression patterns are consistent with their developmental origin, we collected scRNA-seq data from 3 different developmental stages, namely E11.5 to E14.5 (n = 74,550), postnatal day 0 (P0, n = 138,613) and P14 (n = 360,748). These cells were mapped onto the whole mouse brain taxonomy^14,72^ and cells belonging to the subpallium GABAergic cell types were selected. This resulted in a developmental dataset containing 10,259 E11.5 to E14.5 cells, 31,672 P0 cells, and 57,507 P14 cells (**Supplementary Table 4**). This data was integrated with the adult P56 subpallium GABAergic neurons to visualize gene expression patterns across time (**Figure 6b**). From this integration we could delineate which progenitor domain gives rise to specific classes and subclasses in the adult dataset (**Figure 6c,d**).

**Figure 6.**
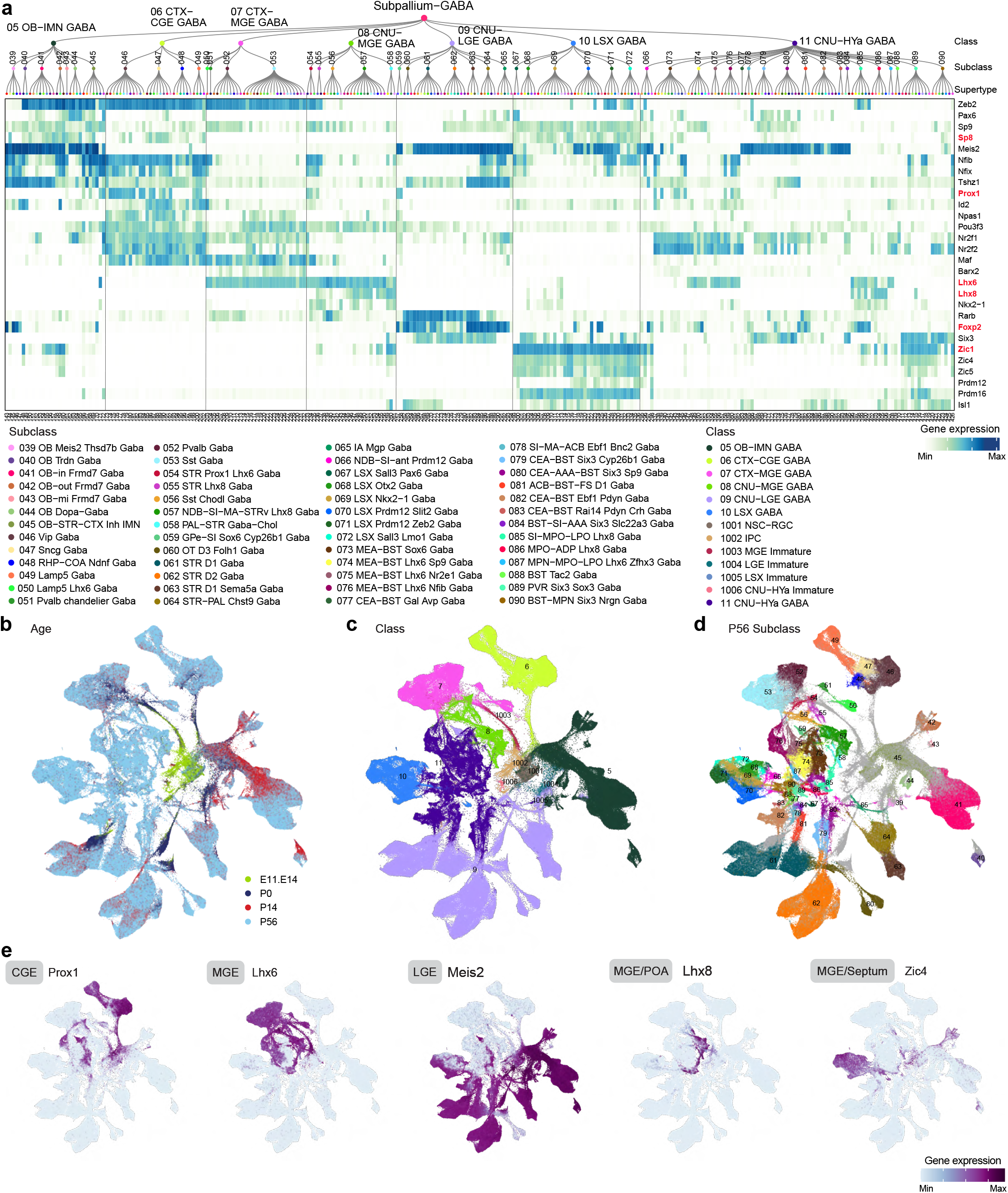
Transcription factor expression in telencephalic GABAergic neurons. **(a)** The supertype dendrogram from Figure 1, followed by a heatmap showing the expression of key transcription factors in each supertype in the taxonomy tree. **(b-e)** UMAP representation of all cell types across the developmental time course, E11.5-E14.5, P0, P14, and P56, colored by age (b), class (c), P56 subclass (d), and major developmental lineage gene markers (e).

#### Developmental CGE-MGE

Starting in early development, CGE- and MGE-derived GABAergic neurons that will populate the CTX and CNU can be identified by their distinct gene expression patterns (**Figure 6e, Extended Data Figure 14a**). The spatial specificity of some transcription factors was verified using the developing mouse ISH atlas^73^ and the P56 MERFISH dataset (**Extended Data Figure 14b**). Our MERFISH dataset was generated using a 500-gene probe set. We imputed the scRNA-seq data into the MERFISH data^14^. By doing so, the imputed spatial expression patten of every gene in the transcriptome, including TF genes not in the MERFISH probe set, can be visualized in the MERFISH sections and confirmed by ISH data^74^ (**Extended Data Figure 14b**).

The cortical CGE and MGE-derived GABAergic classes are predominantly located in all regions of isocortex, OLF, HPF and CTXsp (**Extended Data Figure 1a**), and are marked by expression of the developmental TF *Maf* (**Figure 6a, Extended Data Figure 14a**). The CTX-CGE GABA class specifically expresses the developmental TF *Nr2f2* and *Prox1*. MGE gives rise to GABAergic neurons that populate both cerebral cortex (CTX-MGE GABA class) and striatum and pallidum (CNU-MGE GABA class). Both classes are marked by expression of *Lhx6* and the CNU-MGE GABA class also specifically expresses *Lhx8* (**Extended Data Figure 14b**).

As mentioned above, cortical MGE-derived GABAergic neurons adopt their laminar distribution in an ‘‘inside-out’’ manner that correlates with their birthdate, later-born neurons migrate radially past earlier-born neurons to populate more superficial layers^75,76^. For both CGE and MGE-derived GABAergic neurons we observed an expansion of distinct types that happens between P0 and P14 (**Extended Data Figure 15a-e**). *Sst*-positive GABAergic neurons are among the first types to emerge and diversify, compared to *Pvalb*-positive GABAergic neurons^11,77^. Cortical *Sst*-positive GABAergic neurons arrive at their final destination by P5 in mice but are not yet fully mature at that point^78^. A recent study showed that select subtypes of *Sst*-positive neurons, including *Sst*-positive LRP, are already present in the embryonic cortex but several types of *Sst*-positive interneurons diversify during the postnatal period^68^. *Pvalb*-positive GABAergic neurons exhibit a similar diversification pattern during development. Specifically, the *Pvalb*-positive Chandelier cells show early specification while many other subtypes specify in the early postnatal period^79^. It is becoming increasingly clear that circuit activity influences cell (sub)type specification in cortical GABAergic neurons during the early postnatal period^80,81^.

In corroboration with previous results, the CTX-MGE GABA subclass 50 Lamp5 Lhx6 Gaba expresses both *Lhx6* and *Adarb2*, markers for MGE and CGE respectively (**Extended Data Figure 15f**). As stated above, in the current WMB taxonomy this subclass is assigned to the CTX-MGE class due to expression of the MGE marker *Lhx6*. *Lhx6* and *Adarb2*, marking MGE- and CGE-originated inhibitory neurons respectively, are co-enriched in the POA-derived neurogliaform cells, indicating that the Lamp5 Lhx6 subclass might be POA-derived. From our developmental data here, we can see that at P0 the Lamp5 Lhx6 subclass is more closely related to the CTX-MGE class than the CTX-CGE class, but cluster 10087 has signatures from both CGE and MGE that connects immature CGE and MGE neurons with the Lamp5 Lhx6 subclass (**Extended Data Figure 15b-f**).

#### Developmental LGE

The OB-IMN GABA and CNU-LGE GABA cell classes both arise from the LGE domain (**Figure 6b-e, Extended Data Figure 16a-c**). The OB-IMN GABA class contain cell types located in the MOB and AOB as well as immature neurons in the SVZ lining the lateral ventricles (**Extended Data Figure 1a**). These cells mostly express TFs *Sp8*, *Sp9*, and *Meis2* (**Figure 6e, Extended Data Figure 14a,b**). The types within the CNU-LGE GABA class are predominantly located in STRd and STRv and are marked by expression of *Rarb* and *Foxp2* (**Extended Data Figure 14a,b**).

In the adult GABAergic cell type taxonomy, we observed a relatedness between subclass 39 OB Meis2 Thsd7b Gaba and subclass 65 IA Mgp Gaba from CNU that would suggest that these subclasses share their developmental origin. Subclass 39, containing the neurons that populate the Ipl/Mi, EPl, and Gl layers of OB, is sequestered away from most other olfactory bulb cell types and more closely related to subclass 65 containing immature cells populating the intercalated amygdalar nucleus (**Figure 6d, Extended Data Figure 16b,c)**.

Despite their similarity in adulthood, the distinction between 61 STR D1 Gaba and 62 STR D2 Gaba subclasses is established in early development (**Extended Data Figure 16a-e**). The D1 and D2 populations appear to arise from different progenitor populations based on expression of distinct gene sets (**Extended Data Figure 16f-j**). Genes like *Sp9* and *Six3* are essential for the generation of STR D2 neurons, and genes including *Isl1* and *Ebf1* drive differentiation towards STR D1 neurons (**Extended Data Figure 16f,k-n**). Based on the sparse developmental data collected in this study, it appears that the maturation of the STR D2 neuron population is slower than that of the STR D1 population. At P0 there is less differentiation into mature D2 supertypes compared to D1 supertypes. This delay can also be seen by a set of genes that is expressed at earlier stages in D1 neurons than in D2 neurons (**Extended Data Figure 16e,f,o-q**). In adulthood we observed that STR D1 and STR D2 neuronal types that share the same spatial location are transcriptomically highly similar to each other (**Figure 5**). The transcriptomic distance between STR D1 and STR D2 types is greater during development than in adulthood (**Extended Data Figure 16u**). There are genes in both D1 and D2 types that are expressed in just one of the types at early timepoints but in both types at P14 and P56 (**Extended Data Figure 16f,r-t**). This indicates that the two types develop along their own trajectory but when they reach their final anatomical location their transcriptomes are molded towards their neighboring cells.

#### CNU-HYa and LSX

The LSX GABA class is located specifically in lateral septum complex (LSX) and expresses TFs *Zic1*, *Zic4*, *Zic5*, and *Prdm16* (**Figure 6e, Extended Data Figure 1a, Extended Data Figures 14a,b, 17d**). Lastly, the CNU-HYa GABA class contains cell types present in both CNU, mostly located in sAMY, PALc, and POA. Cell types in LSX and CNU-HYa are highly diverse and have multiple embryonic origins. LSX contains a population of *Nkx2-1*-positive neurons that might arise from MGE or POA. CNU-HYa cell types located in CEA, AAA, and BST express *Meis2* and *Six3*, cell types in MEA and certain BST cell types express *Lhx6* and *Nr2f2*, and cell types located in POA express *Zic1* and *Zic4* (**Extended Data Figure 17d**). From the sparse developmental data we collected it was difficult to precisely link the developmental origin to each adult subclass (**Extended Data Figure 17a-c**). Both LSX and CNU-HYa classes show similar progenitor populations at E11.5-E14.5, which are very distinct from the cell populations detected at P0. While there is a great expansion of types between P0 and P14 in the cortical GABAergic cell types, in the LSX GABA and CNU-HYa GABA classes this is not the case. At P0, we identified nearly all adult cell types within these classes, indicating that cell type diversification completed before birth similar to what has been described for POA derived hypothalamic neurons^82^.

## DISCUSSION

In this study, we present an extraordinarily complex, high-resolution cell type taxonomy and atlas of GABAergic neurons in the mouse telencephalon. We have integrated high-quality single cell RNA-seq data with an adult whole brain MERFISH dataset. This integration allowed us to analyze the molecular and anatomical organization of GABAergic neurons in detail. We find that telencephalic GABAergic cell types can be organized in a hierarchical manner where relationships among cell types reflect both spatial location and developmental origin. We systematically relate cell types in our taxonomy to the wide variety of previously identified cell types or populations across all telencephalic regions, and we discover and categorize many new cell types that were previously unknown or with little information. As such, the telencephalic GABAergic cell type taxonomy can serve as a foundational reference for molecular, structural and functional studies of cell types and circuits by the entire community. Furthermore, we gain additional insights into the organization of these cell types, including multiple dimensions of continuous variations among related cell types along with correlated gene expression gradients, long-range migration and dispersion to distinct brain regions of closely related cell types with a common developmental origin, and a stark contrast in pre- and postnatal diversification between cortical/striatal/some pallidal and septal/other pallidal/preoptic GABAergic cell types.

Telencephalic GABAergic neurons originate from five principal progenitor domains of the subpallium: MGE, CGE, LGE, the embryonic POA, and the embryonic septum^3,5,83^. From here the progenitor cells migrate to distant locations and differentiate to form mature brain structures. These processes are regulated by morphogen-regulated transcription factor modules. Our understanding of the TF cascades that specify neuronal cell types during development has increased greatly^11,83–85^. Many of the TFs regulating neuronal differentiation during development are also expressed in mature neurons and we can use the expression of these TFs to infer the expected developmental origin of neuronal subtypes. More recently, studies have shown that some of these TFs also play a role in the maintenance of neuronal identity. In invertebrates, the term “terminal selector gene” has been used to describe genes that not only specify but also maintain cell type identity^86,87^ and these terminal selectors often work in specific combinations to define neuronal subtypes. In mouse, the term “master regulator” has been used to describe the transcription factor(s) that triggers the gene expression program of a developmental lineage. Many of these TFs have been identified by the fact that genetic removal during development results in failure of specific neuronal classes to develop properly. Studies on the role of these regulators in maintenance of neuronal identity are limited but there is reason to believe these terminal selector genes exist in mice as well^88,89^. Our study reveals a large set of persistent TFs in the telencephalic GABAergic cell types (**Figure 6**), and their roles in the maintenance of cell type identity can be tested in future genetic perturbation experiments.

A nearly universal characteristic of the telencephalic GABAergic neurons is that neurons located in far apart regions of the telencephalon can belong to the same cell type (subclass, supertype, or even cluster). This suggests that neurons of a common developmental origin can migrate long distances and reside in highly distinct brain regions. Most supertypes and clusters in the CTX-CGE and CTX-MGE classes, that is, the CGE- and MGE-derived cortical GABAergic interneurons, are widely distributed in nearly all isocortex, HPF, OLF and CTXsp regions, with only a small group of supertypes and clusters located selectively in HPF, OLF and/or CTXsp (**Figure 2, Extended Data Figure 5**). The CNU-MGE class, containing all striatal GABAergic interneurons and many pallidal GABAergic neurons, shares transcriptomic similarity and a common origin from MGE^90–92^ to the CTX-MGE class which includes the cortical *Pvalb*-positive and *Sst*-positive GABAergic interneurons (**Figure 2**). The OB-IMN class, containing olfactory bulb GABAergic neurons, developmentally originates in LGE and is transcriptomically related to the CNU-LGE class which contains the striatal D1 and D2 MSNs and related cell types (**Figure 3, Extended Data Figures 3, 16**). The OB-IMN class also contains immature neurons generated in SVZ through adult neurogenesis and migrating to OB. Lastly, the CNU-HYa class contains a large set of highly heterogeneous cell types that are widely distributed in sAMY, medial, ventral and caudal pallium, as well as the hypothalamic preoptic area (**Figure 4, Extended Data Figure 9**).

At more fine-grained levels, subclass 56 Sst Chodl Gaba in the CNU-MGE class contains both the cortical Sst-Chodl neurons (as a specific supertype) that are the long-range projection neurons enriched in deep layers of many cortical areas^28,67,93^, and the *Sst*-positive interneurons in the striatum^32,94^ (also found in pallidal regions) that have electrophysiological properties similar to that of MSNs^32,95^ (**Figure 2**). As such, the cortical Sst-Chodl neurons are more similar to the striatal/pallidal Sst -Chodl interneurons than the cortical Sst interneurons. The cholinergic neurons in the 58 PAL-STR Gaba-Chol subclass, also belonging to the CNU-MGE class, include the basal forebrain cholinergic projection neurons in multiple pallidal regions as well as the striatal cholinergic interneurons, with spatial specificity at supertype or cluster level (**Figure 2, Extended Data Figure 11**). Lastly, within multiple subclasses in the CNU-HYa class, neurons belonging to the same supertype can be found in both CEA (part of sAMY) and BST (part of PALc), or in both MEA (part of sAMY) and BST (**Figure 4, Extended Data Figure 9**).

As a consequence of this widespread migration and dispersion, most of the telencephalic regions (except for LSX and OB) contain a heterogeneous mixture of GABAergic neuronal types (**Extended Data Figure 1**) with distinct developmental origins and likely distinct connectivity and circuit functions. Within the major developmental progenitor domains, several functional subdomains have been identified that generate different types of GABAergic neurons. For example, the LGE subventricular zone can be divided into four subdomains along the dorsoventral axis (pLGE1, 2, 3, and 4). The dorsal domain (dLGE; pLGE1, 2) expresses genes like *Sp8* and *Er81* and gives rise to olfactory bulb interneurons and intercalated cells (ITCs) of the amygdala, and the ventral domain (vLGE; pLGE3, 4) expresses genes like *Isl1* and *Ebf1* predominantly gives rise to GABAergic medium spiny neurons^38,96,97^. More recent studies indicate that the pLGE3 and pLGE4 domains might preferentially generate MSN D2 and MSN D1 neurons respectively^98,99^. Similarly, MGE can be divided into five subdomains along the dorsoventral axis (pMGE1 to 5) and each of these domains generates multiple types of GABAergic neurons^5,100,101^. The more ventral part of MGE (pMGE4,5) tends to generate many GABAergic neurons that populate the striatum and pallidum, while the dorsal part of MGE (pMGE1-4) generates many of the cortical interneurons^5,101,102^. Future studies will be needed to systematically understand the spatial and temporal patterns of the emergence of diverse cell types within each progenitor domain.

As the GABAergic neurons reach their final destinations in later stages of development, cell type identities may be further shaped by local environment, as evidenced by gene expression gradients and continuous variations in various spatial dimensions across supertypes and clusters of a specific GABAergic subclass. The MGE-derived cortical *Sst*- and *Pvalb*-neurons exhibit continuous variations from deep to superficial layers (**Extended Data Figure 12**). The six LSX subclasses are not spatially segregated but partially overlapping, and they exhibit multidimensional gradients (**Extended Data Figure 13**) that might align with their input/output connections^69,70^. The most prominent continuous variations we observed are in the striatal D1 and D2 MSN types (**Figure 5**). We have identified 5 subclasses of MSN types of which subclasses 61 STR D1 Gaba and 62 STR D2 Gaba contain the “classical” MSN D1 and MSN D2 neurons that are present in striatum and have defined gene expression profiles, connectivity patterns, and function^33,34,103–106^. These two subclasses show a strong correlation along the dorsolateral-to-ventromedial axis of striatum (**Figure 5**). We have identified gene modules, shared between MSN D1 and MSN D2 types, that drive the gene expression gradient along the dorsolateral-to-ventromedial axis of striatum. Genes involved in cGMP-PKG signaling are expressed at higher levels in the dorsolateral MSN D1 and D2 types. Downstream effectors of this signaling pathway play a key role in regulating long-term changes in striatal synaptic efficacy^107–109^. On the ventromedial end of the gradient axis genes coding for various neuroactive receptors are expressed at higher levels of which *Cnr1* has been described before ^34,110^. The dorsolateral-to-ventromedial transcriptional gradient and cell types related to that align with topographical organization of excitatory striatal afferent projections^111,112^.

We have identified distinct cholinergic neuronal types in striatum, pallidum, and lateral septum. Most telencephalic cholinergic neurons originate from the MGE, embryonic POA, and embryonic septum^113,114^. The cholinergic precursors originate from the *Nkx2.1*-positive domain and are further specified by combinatorial expression of additional transcription factors^5^. *Lhx6* is essential for specification and migration of MGE-derived GABAergic interneurons, in both the cortex and striatum^6,115,116^ and *Lhx8* has been associated with the specification of a cholinergic phenotype by actively inducing cholinergic properties^6^. These genes are still expressed in the cholinergic neurons in adulthood (**Extended Data Figure 11e**). The cholinergic neurons acquire different identities based on their time of birth, which occurs between E12 and E16^117^. Early- and late-born cholinergic striatal interneurons migrate at different time points and populate different regions. Absence of *Gbx2* has been shown to cause ablation of the entire late-born cholinergic population^118^. This late-born population most likely corresponds to supertype 260 PAL-STR Gaba-Chol_2 which still expresses *Gbx2* in adulthood and is located in striatum. A subset of the striatal cholinergic interneurons in this supertype expresses the Type-3 vesicular glutamate transporter (*VgluT3*, *Slc17a8*; **Extended Data Figure 11e**) and can mediate glutamatergic transmission which is required for cholinergic signaling onto fast spiking interneurons (subclass 54 STR Prox1 Lhx6 Gaba) as well as acetylcholine-dependent inhibition of MSNs^119,120^. While striatal cholinergic neurons mostly serve as local interneurons, pallidal cholinergic neurons are mostly sending projections to cortex, hippocampus, and amygdala. These basal forebrain cholinergic neurons are distributed across a series of nuclei, including MS, NDB, the nucleus basalis of Meynert (part of SI), and GPe. Several studies have shown that the function of basal forebrain cholinergic neurons is linked to their topographical organization. For example, the dorsal parts of PL, ILA, and ACA of isocortex receive projections from medially located SI and NDB neurons, whereas more ventral parts of prefrontal cortex receive projections from more laterally located basal forebrain nuclei. Hippocampus and the entorhinal cortex receive the majority of their cholinergic input from the MS and vNDB cholinergic neurons^121,122^.

Finally, the developmental scRNA-seq data presented in this study reveals an interesting difference between the MGE/CGE/LGE derived GABAergic neuronal types (in classes CTX-CGE, CTX-MGE, CNU-MGE, CNU-LGE and OB-IMN) and those in the LSX and CNU-HYa GABAergic classes. The former group of GABAergic types exhibit relatively clear continuity from E11-14 to P0 to P14 to P56, along with substantial transcriptomic shifts between E11-14 and P0 as well as between P0 and P14 (**Extended Data Figures 15, 16**), suggesting these cell types undergo continued and extensive postmitotic and postnatal diversification similar to what has been described before for cell types in cortex^123,124^. On the other hand, the latter group of septal and most pallidal GABAergic types exhibit disjointed transcriptomic changes between E11-14 and P0, whereas no substantial transcriptomic shifts from P0 to P14 to P56 were observed (**Extended Data Figure 17**), suggesting that this cell type repertoire emerges in an apparent burst in the embryonic stage with limited postnatal diversification, consistent with a recent study^82^. This developmental difference between these two groups of GABAergic neuronal types is intriguing because it is consistent with our previous observation of the dichotomy of cell type characteristics between the dorsal and ventral parts of the adult brain^14^.

In conclusion, our study provides a detailed transcriptomic characterization of GABAergic neurons in the telencephalon, their spatial locations, and their potential developmental origins. It highlights both the vast differences and the similarities between spatially distant and not so distant types of GABAergic neurons. With the current developmental dataset, we could link adult cell types to their developmental origins, but more detailed molecular investigations will be needed to fully understand how neurons across different lineages diversify during development. Moreover, though the spatial organization of transcriptomic types follows a similar logic to the current knowledge of the circuit organization in the telencephalon, further experiments are needed to link the transcriptomic types to their projection and connectivity patterns.

## METHODS

### Sample collection, data generation, and data analysis for P56 dataset

Most of methods which apply to the adult P56 10x scRNA and MERFISH datasets used for this paper were described before^14^ and the following methods are either newly introduced or modified version for this paper.

### UMAP projection

We performed PCA based on the imputed gene expression matrix of 4,895 marker genes using the 10xv3 reference. We selected the top 100 PCs, then removing one PC with more than 0.7 correlation with the technical bias vector, defined as log2(gene count) for each cell. We used the remaining PCs as input to create 2D and 3D UMAPs^125^, using parameters nn.neighbors = 25 and md = 0.4.

### Constellation plot

To generate the constellation plot, each transcriptomic supertype was represented by a node (circle), whose surface area reflected the number of cells within the supertype in log scale. The position of nodes was based on the centroid positions of the corresponding supertypes in UMAP coordinates. The relationships between nodes were indicated by edges that were calculated as follows. For each cell, 15 nearest neighbors in reduced dimension space were determined and summarized by supertypes. For each supertype, we then calculated the fraction of nearest neighbors that were assigned to other supertypes. The edges connected two nodes in which at least one of the nodes had > 5% of nearest neighbors in the connecting node. The width of the edge at the node reflected the fraction of nearest neighbors that were assigned to the connecting node and was scaled to node size. For all nodes in the plot, we then determined the maximum fraction of “outside” neighbors and set this as edge width = 100% of node width. The function for creating these plots, plot_constellation, is included in scrattch.bigcat.

### Imputation of scRNA-seq data into the MERFISH space

The MERFISH dataset was collected using only 500 genes. To obtain the spatial distribution of all the genes, we projected gene expression of the MERFISH dataset to the 10xv3 scRNA-seq dataset using a modified version of the impute_knn_global function in the scrattch.bigcat package^14,26^. We used self-imputed 10xv3 dataset as reference, meaning that the expression of each 10xv3 cells was first imputed based on its nearest 15 neighbors in the reduced principal component space. This decision was made to ensure that in the following hierarchical imputation step, the transitions between major cell types were preserved. The imputation was conducted in the order specified by the hierarchy defined by class and subclass. At the root, we imputed the expression for all the genes for each MERFISH cell based on the average expression of their nearest neighbors from the reference 10xv3 dataset, defined by the cosine similarity using all 500 MERFISH genes. In each of the following iterations, we selected the node to which each MERFISH cell was assigned and imputed only the expression of the DEGs based on pairwise comparison for all the clusters under this node. The nearest neighbors for imputation were selected from the clusters under this node in the reference dataset, using only the subset of DEGs that were present on the MERFISH gene panel. We repeated this procedure until reaching the leaf node. This strategy enabled us to preserve the cell type resolution during imputation, making it less susceptible to the global platform differences between MERFISH and scRNA-seq.

### Analysis of spatial gene expression gradients

We performed independent component analysis using fastICA^126^ to decompose the gene expression matrix into independent components. These components are then projected onto the imputed MERFISH data to determine if the component represents a spatial gene expression program. For components that represent a spatial gene expression program the top loading genes we selected and visualized on both the UMAP in RNAseq space and on sections in the imputed MERFISH space. We evaluated the gene modules in the identified individual components and applied UCell^127^ to assign a “gene module score” based on both positive and negative genes to each cell.

### Spatial domain clustering

We used BANKSY^128^ to perform spatial domain clustering within BST neurons. This algorithm implements a feature-augmentation approach to map domains by integrating the transcriptional profiles of individual cells with their physical distances and tissue neighborhood context. As the MERFISH data was registered to the CCFv3, it allowed us to subset BST neurons from the MERFISH data. We used the 8988 BST neurons, their spatial location, and their 500 genes expression profile as input for BANKSY.

### Assessing concordance of cell type taxomony between the subpallium GABAergic cell type atlas and external datasets

We performed mapping of cells from each external dataset to the 10x v3 whole-brain dataset using treeMap function from scrattch.mapping package (https://github.com/AllenInstitute/scrattch-mapping)^72^. The reference cell type taxonomy was organized in a hierarchy defined by class and subclass. At each node, top markers were selected that best discriminate the clusters belonging to different child nodes. Starting at the root, cells were assigned to the closest cluster centroid from all the clusters under the given node based on the selected node markers using the cosine similarity metric. This mapping procedure was repeated until reaching the leaf nodes. To assess mapping confidence, we subsampled 80% of the markers at each node, and repeated the mapping process 100 times. In each bootstrapping step, we also computed the cosine similarity of the cell to the mapped the cluster based on the markers for all the nodes along the mapping path and calculated the average similarity across all 100 bootstrapping iterations. This score was used to assess the quality of the mapping. Cells with a score above 0.5 were used to generate a confusion matrix showing the proportion of cells jointly found between 2 types and their Jaccard similarity score.

### Measuring similarity between MSN D1 and D2 clusters

We computed the nearest neighbors from MSN D1 cells for MSN D2 cells and vice versa using cosine similarity metric based on the same marker list, which were used to define the edge weights for the constellation plots. To select the most similar pairs between MSN D1 and MSN D2 types, we selected all the pairs with Pearson correlation greater than 0.93 and with at least 30% of the KNNs from one cluster belonging to the other cluster in the pair.

### Developmental scRNA-seq data collection

#### Mouse breeding and husbandry

All experimental procedures related to the use of mice were approved by the Institutional Animal Care and Use Committee of the AIBS, in accordance with NIH guidelines. Mice were housed in a room with temperature (21–22 °C) and humidity (40–51%) control within the vivarium of the AIBS at no more than five adult animals of the same sex per cage. Mice were provided food and water ad libitum and were maintained on a regular 14:10 h light:dark cycle. Mice were maintained on the C57BL/6 J background. We excluded any mice with anophthalmia or microphthalmia.

Mothers of all experimental pups were placed in a fresh cage when embryos were ∼E8. We used 6 pups to collect 74,550 cells from ages E11.5, E12.5, E13.5, and E14.5. From ages E11.5 and E12.5 we collected whole brain tissue and from ages E13.5 and E14.5 we collected cerebrum and brain stem (CH-BS). From 6 P0 pups we collected 138,613 cells, and from 6 P14 pups we collected 360,748 cells. P0 and P14 cells were collected from both male and female mice across 6 dissection ROIs: OLF, CTXsp, Isocortex, HPF, CNU, and HY. No statistical methods were used to predetermine sample size. All donor animals used for the developmental scRNA-seq data generation are listed in **Supplementary Table 5**. Brain dissections for all groups took place in the morning.

#### Single-cell isolation

Single cells were isolated following a cell-isolation protocol developed at AIBS^129^. The brain was dissected, submerged in artificial cerebrospinal fluid (ACSF), embedded in 2% agarose, and sliced into 350-μm coronal sections on a compresstome (Precisionary Instruments). Block-face images were captured during slicing. ROIs were then microdissected from the slices and dissociated into single cells.

Dissected tissue pieces were digested with 30 U ml−1 papain (Worthington PAP2) in ACSF for 30 min at 30 °C. Due to the short incubation period in a dry oven, we set the oven temperature to 35 °C to compensate for the indirect heat exchange, with a target solution temperature of 30 °C. Enzymatic digestion was quenched by exchanging the papain solution three times with quenching buffer (ACSF with 1% FBS and 0.2% BSA). Samples were incubated on ice for 5 min before trituration. The tissue pieces in the quenching buffer were triturated through a fire-polished pipette with 600-µm diameter opening approximately 20 times. The tissue pieces were allowed to settle and the supernatant, which now contained suspended single cells, was transferred to a new tube. Fresh quenching buffer was added to the settled tissue pieces, and trituration and supernatant transfer were repeated using 300-µm and 150-µm fire-polished pipettes. The single-cell suspension was passed through a 70-µm filter into a 15-ml conical tube with 500 µl of high-BSA buffer (ACSF with 1% FBS and 1% BSA) at the bottom to help cushion the cells during centrifugation at 100g in a swinging-bucket centrifuge for 10 min. The supernatant was discarded, and the cell pellet was resuspended in the quenching buffer. The concentration of the resuspended cells was quantified, and cells were immediately loaded onto the 10x Genomics Chromium controller.

#### cDNA amplification and library construction

The E11.5 to E14.5 cell suspensions were processed using the Chromium Single Cell 3′ Reagent Kit v3 (1000075, 10x Genomics). We followed the manufacturer’s instructions for cell capture, barcoding, reverse transcription, cDNA amplification and library construction^130^. We loaded 8,283 ± 703 cells per port. We targeted a sequencing depth of 120,000 reads per cell; the actual average achieved was 70,324 ± 62,149 reads per cell across 9 libraries.

The P0 cell suspensions were processed using the Chromium Single Cell 3′ Reagent Kit v3.1 (1000268, 10x Genomics). We followed the manufacturer’s instructions for cell capture, barcoding, reverse transcription, cDNA amplification, and library construction^131^. We loaded 11,551 ± 1,785 (mean ± s.d.) cells per port. We targeted sequencing depth of 120,000 reads per cell; the actual average achieved was 65,069 ± 61,474 (mean ± s.d.) reads per cell across 12 libraries.

The P14 cell suspensions were processed using the Chromium Next GEM Single Cell 3’ HT Reagent Kit v3.1 (1000370, 10x Genomics). We followed the manufacturer’s instructions for cell capture, barcoding, reverse transcription, cDNA amplification, and library construction. We loaded 30,062 ± 15,008 (mean ± s.d.) cells per port. We targeted sequencing depth of 120,000 reads per cell; the actual average achieved was 46,055 ± 61,941 (mean ± s.d.) reads per cell across 12 libraries.

#### Sequencing data processing and QC

Processing of 10x Genomics scRNA-seq libraries was performed as described previously^26^. In brief, libraries were sequenced on the Illumina NovaSeq6000, and sequencing reads were aligned to the mouse reference transcriptome (M21, GRCm38.p6) using the 10x Genomics CellRanger pipeline (version 6.1.1) with default parameters. To remove low-quality cells, we applied similar QC analysis and thresholding as described previously^14^.

#### Clustering scRNA-seq data

To assign cell type identity to cells at P14 the cells were mapped onto the WMB taxonomy using Hierarchical Approximate Nearest Neighbour (HANN) mapping available in scrattch-mapping package^72^. To improve mapping to the correct lineage, we removed three classes containing immature neurons (03 OB-CR Glut, 04 DG-IMN Glut, and 05 OB-IMN GABA) from the adult WMB taxonomy. For cells from P0 time point, we assigned their broad cell types by mapping to the nearest cluster centroid in the adjacent older age group, P14, using scrattch.mapping. Cells from E11.5 to E14.5 were binned, considered as one time point for further analysis, and mapped to the P0 time point. After assigning the broad cell types, iterative clustering was performed within assigned subclasses using the scrattch.bigcat package as described before^14^. Cell type annotation of the developmental scRNA-seq dataset is shown in **Supplementary Table 4**.

## ACKNOWLEDGEMENTS

We are grateful to the Transgenic Colony Management, Lab Animal Services, Molecular Biology and Histology teams at the Allen Institute for technical support. The research was funded by the U19MH114830 and U01MH130962 grants from National Institute of Mental Health to H.Z., under the BRAIN Initiative of National Institutes of Health (NIH). The content is solely the responsibility of the authors and does not necessarily represent the official views of NIH and its subsidiary institutes. This work was also supported by the Allen Institute for Brain Science. The authors thank the Allen Institute founder, Paul G. Allen, for his vision, encouragement, and support.

## Author Contributions

Conceptualization: H.Z. Data analysis lead and coordination: C.T.J.vV. Data generation scRNA-seq: C.T.J.v.V., T.C., M.C., R.F., J.Gl., J.Gu., C.R.H, W.H., K.J., R.MC., T.H.P., K.R., E.D.T., A.T., N.D., K.A.S., Z.Y., H.Z. Data processing and analysis (scRNA-seq): C.T.J.v.V., Y.G., C.L., A.B.C., R.C., T.D., J.Go., B.N., K.A.S., Z.Y., H.Z. Data generation (MERFISH): M.K., D.M., J.W., H.Z. Data processing and analysis (MERFISH): C.T.J.v.V., M.K., D.M., S.D., M.H., L.N., J.W., Z.Y., H.Z. Project management: C.P., L.K., K.S. Management and supervision: C.T.J.v.V., D.M., N.D., L.N., J.W., K.A.S., B.T., Z.Y., H.Z. Manuscript writing and figure generation: C.T.J.vV., Z.Y., H.Z. Manuscript review and editing: C.T.J.v.V., M.K., Z.Y., H.Z.

## Additional Information

Correspondence and requests for materials should be addressed to: H.Z. or C.T.J.vV.

## Data Availability

The scRNA-seq and MERFISH datasets for this study are part of the Allen whole mouse brain (WMB) cell type atlas and are accessible through Neuroscience Multi-omic Data Archive (NeMO, https://nemoarchive.org/) and Brain Image Library (BIL, https://www.brainimagelibrary.org/index.html). The 10x scRNA-seq datasets (FASTQ files) are available at NeMO under identifier https://assets.nemoarchive.org/dat-qg7n1b0. The MERFISH dataset is available at BIL under DOI https://doi.org/10.35077/g.610.

The Subpallium-GABA cell type taxonomy is available from the Allen Brain Cell (ABC) Atlas at https://portal.brain-map.org/atlases-and-data/bkp/abc-atlas, to visualize both sc/snRNA-seq and MERFISH datasets. Instruction for access of the processed 10x scRNA-seq data is available at https://github.com/AllenInstitute/abc_atlas_access/blob/main/descriptions/WMB-10X.md, and instruction for access of the processed MERFISH data is available at https://github.com/AllenInstitute/abc_atlas_access/blob/main/descriptions/MERFISH-C57BL6J-638850.md.

The developmental scRNA-seq datasets are being made available through BRAIN Initiative Cell Atlas Network (BICAN), www.portal.brain-bican.org, and at NeMO.

## Code Availability

The R package scrattch.bigcat, is available via github https://github.com/AllenInstitute/scrattch.bigcat. Notebooks with examples of data analysis code used in this manuscript are available via github https://alleninstitute.github.io/scrattch.example/.

---

**Extended Data Figure 1.**
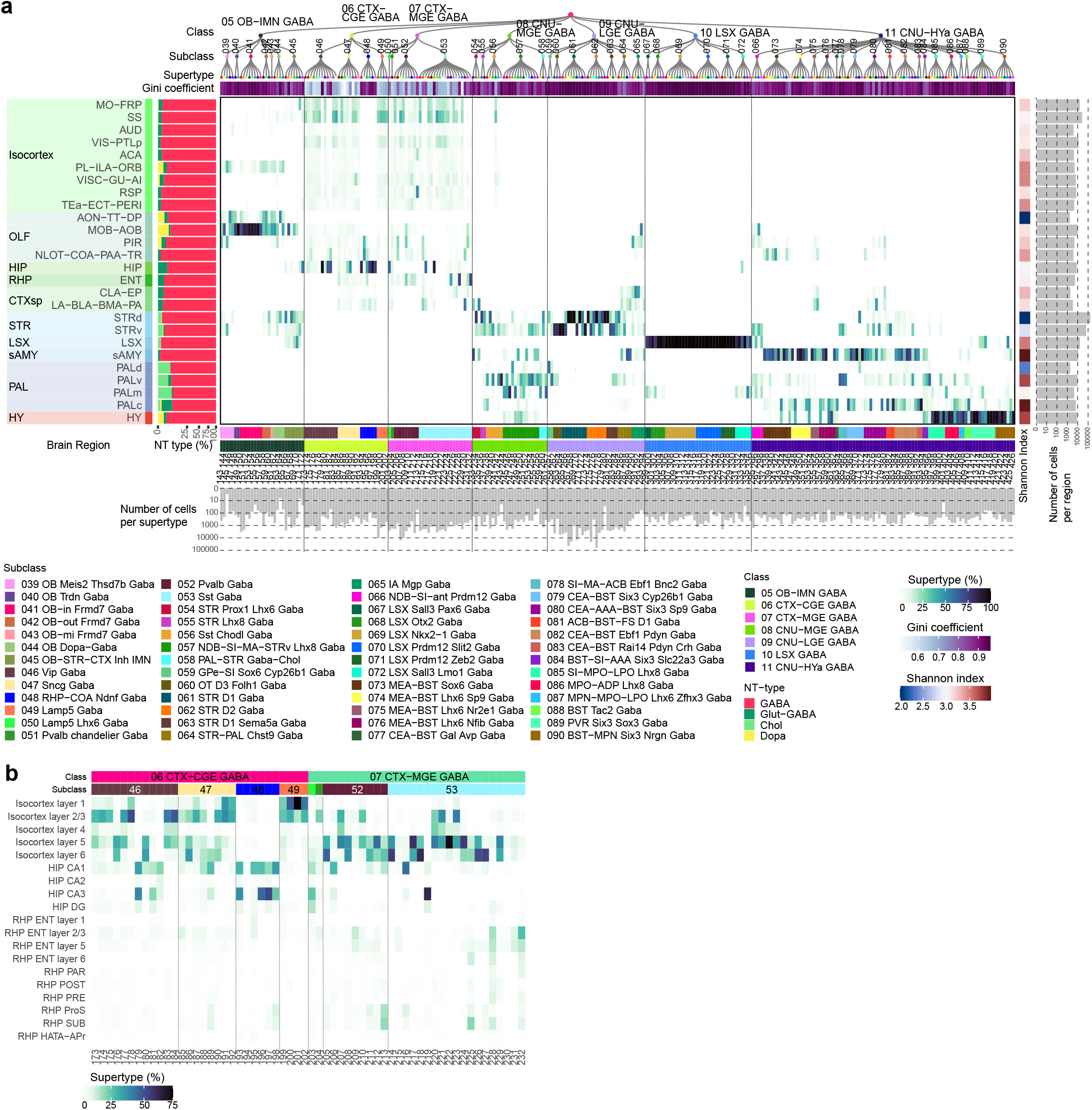
GABAergic neuronal type composition in different regions of the telencephalon. **(a)** Heatmap showing the proportion of cells in each broad region of the telencephalon per GABAergic supertype. (**b**) Heatmap showing the proportion of cells in each supertype from 06 CTX-CGE GABA and 07 CTX-MGE GABA classes in each cortical layer and substructure of the hippocampal formation.

**Extended Data Figure 2.**
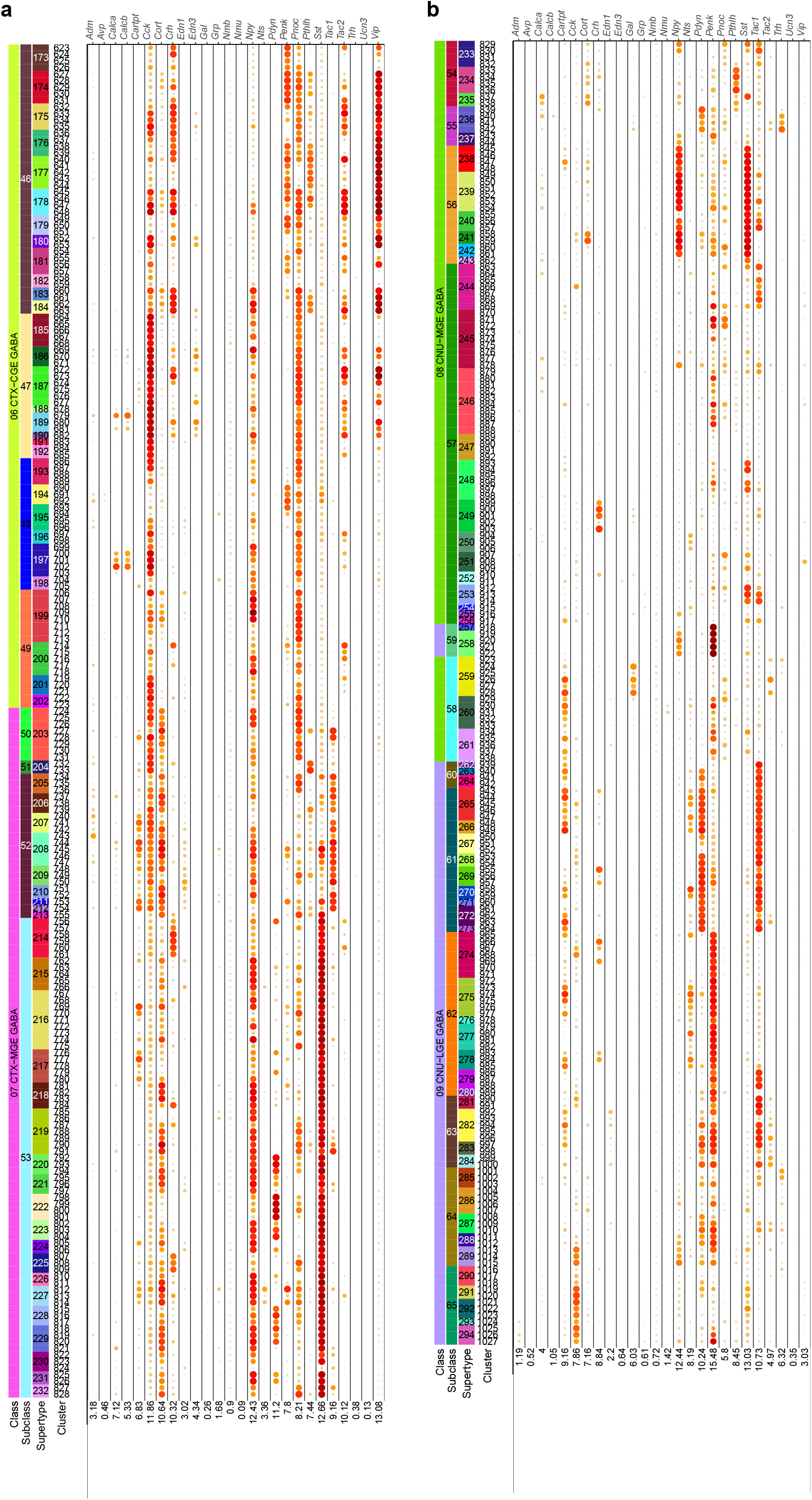
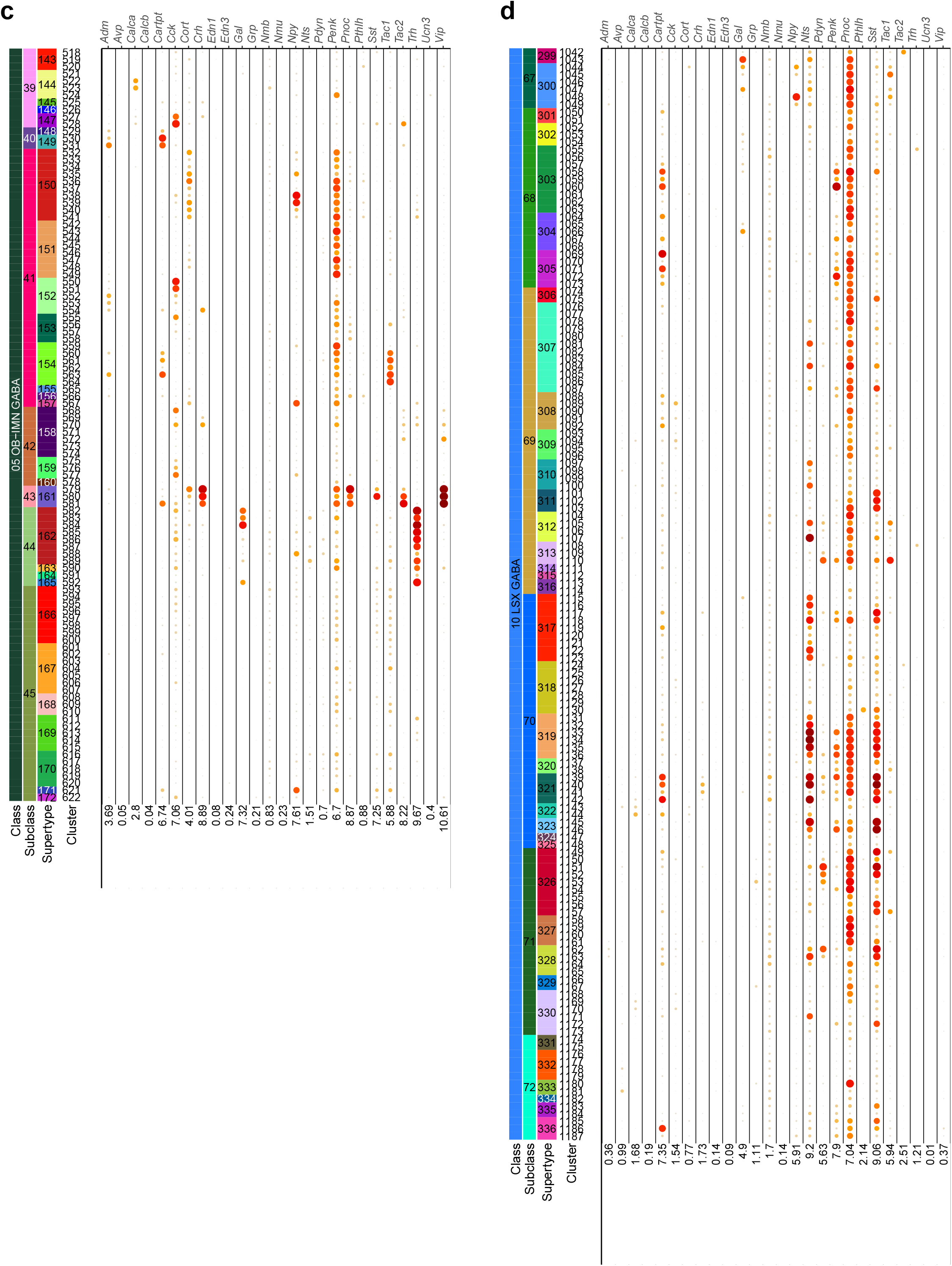
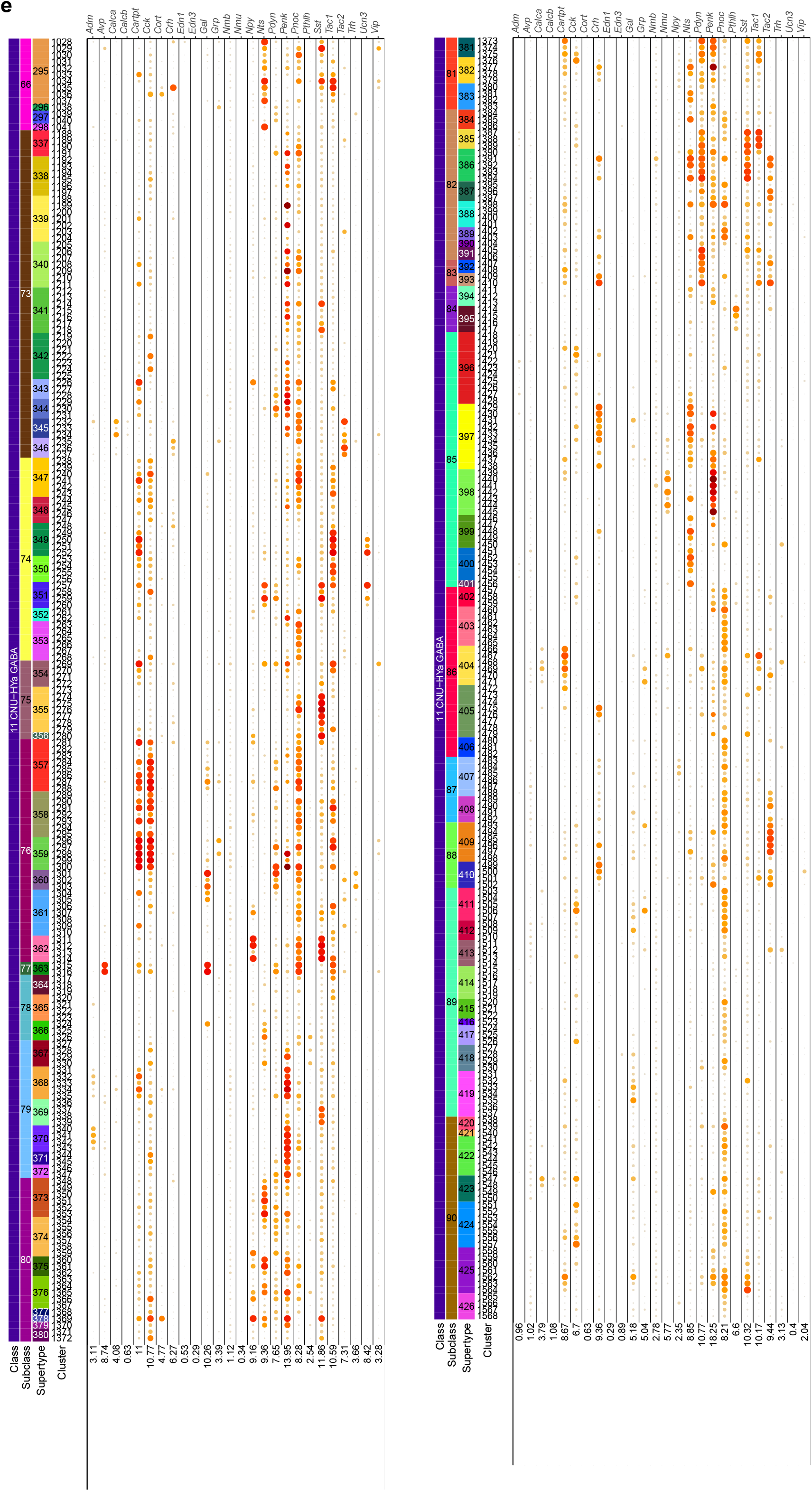
Neuropeptide gene expression in telencephalic GABAergic neuronal types. **(a-e)** Dot plot showing gene expression level of differentially expressed neuropeptides in each cluster across classes 6 CTX-CGE GABA and 7 CTX-MGE GABA (a), 8 CNU-MGE GABA and 9 CNU-LGE GABA (b), 5 OB-IMN GABA (c), 10 LSX GABA (d), and 11 CNU-HYa GABA (e). Dot size and color indicate proportion of expressing cells and average expression level in each cluster, respectively.

**Extended Data Figure 3.**
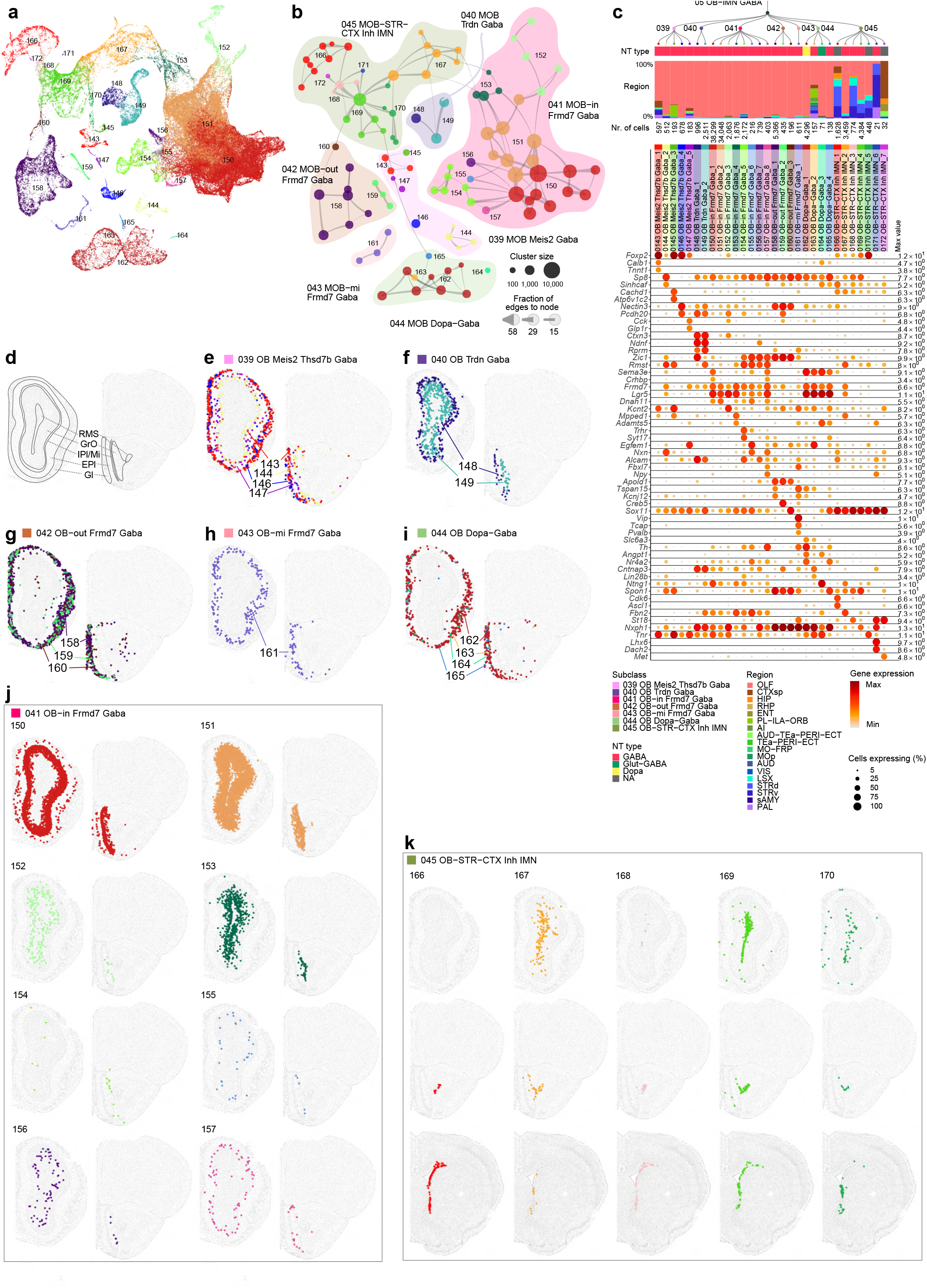
GABAergic and immature neuronal cell types of olfactory bulbs. UMAP representation of GABAergic and immature neuronal types in olfactory bulbs colored by supertype **(a)**. **(b)** Constellation plot of OB-IMN GABA clusters using UMAP coordinates shown in b. Nodes are colored by supertype and grouped in bubbles by subclass. **(c)** Dendrogram of OB-IMN GABA supertypes followed by bar graphs showing major neurotransmitter type and region distribution of profiled cells, followed by dot plot showing marker gene expression in each supertype from the 10xv3 dataset. Dot size and color indicate proportion of expressing cells and average expression level in each supertype, respectively. **(d)** Schematic drawing of anatomical structure in MOB (left) and AOB (right). Abbreviations: RMS, rostral migratory stream; GrO, granular layer; IPl, internal plexiform layer; Mi, mitral layer; EPl, external plexiform layer; Gl, glomerular layer. **(e-k)** Representative MERFISH sections showing the location of OB subclasses 39 MOB Meis2 Gaba (e), 40 OB Trdn Gaba (f), 42 OB-out Frmd7 Gaba (g), 43 OB-mi Frmd7 Gaba (h), 44 OB Dopa-Gaba (i), 41OB-in Frmd7 Gaba (j), and 45 OB-STR-CTX Inh IMN (k). Cells are colored and labelled by supertype.

**Extended Data Figure 4.**
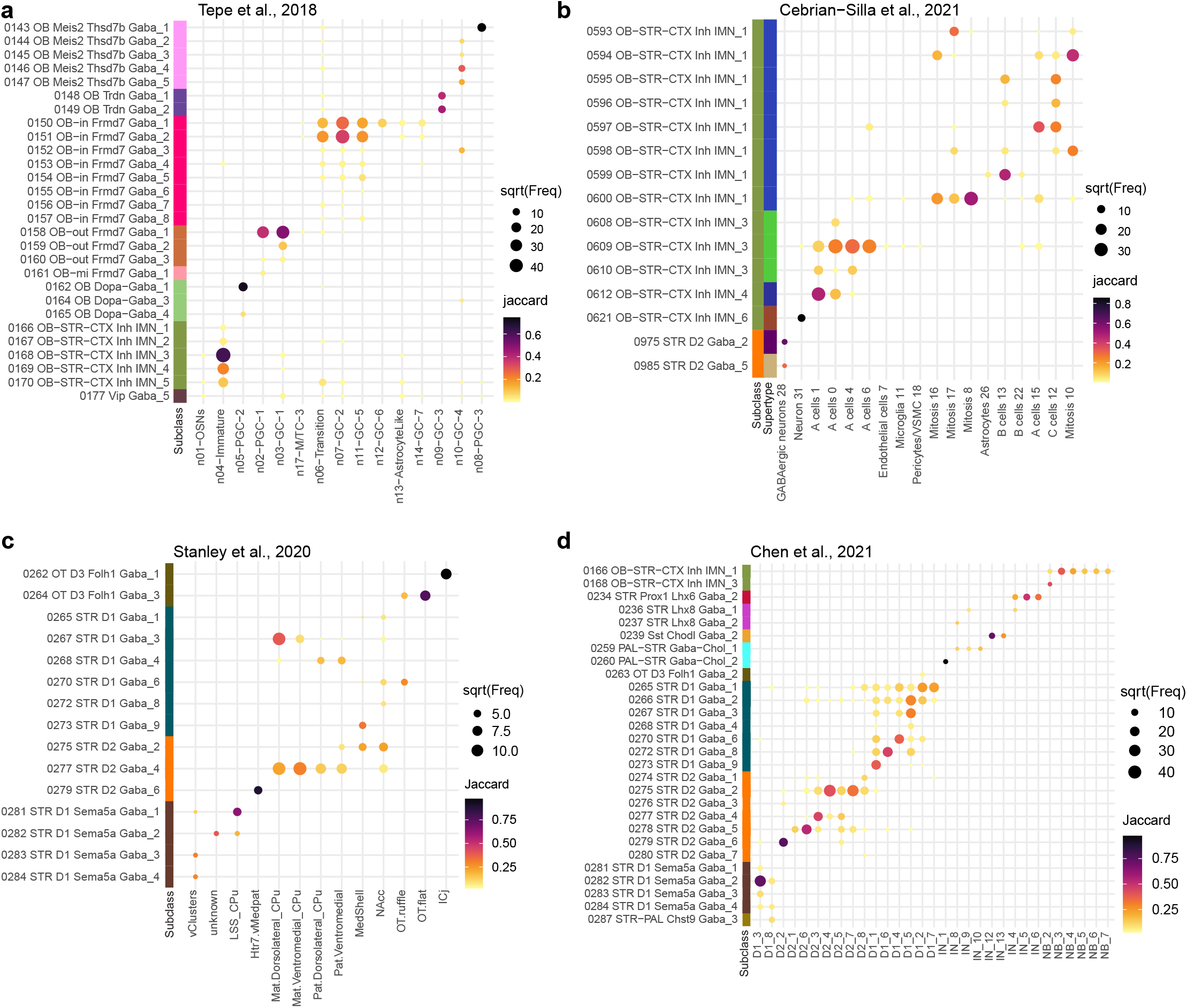
Correspondence between the current transcriptomic taxonomy of OB-IMN GABA and CNU-LGE GABA classes and previously published ones. Correspondence was determined by mapping cells from previously published datasets to the current taxonomy as described before^26^. **(a,b)** Mapping of Tepe et al., 2018^21^ (a) and Cebrian-Silla et al., 2021^18^ (b) to the OB-IMN GABA class. **(c,d)** Mapping of Stanley et al., 2020^34^ (c) and Chen et al., 2021^33^ (d) to the CNU-LGE GABA class.

**Extended Data Figure 5.**
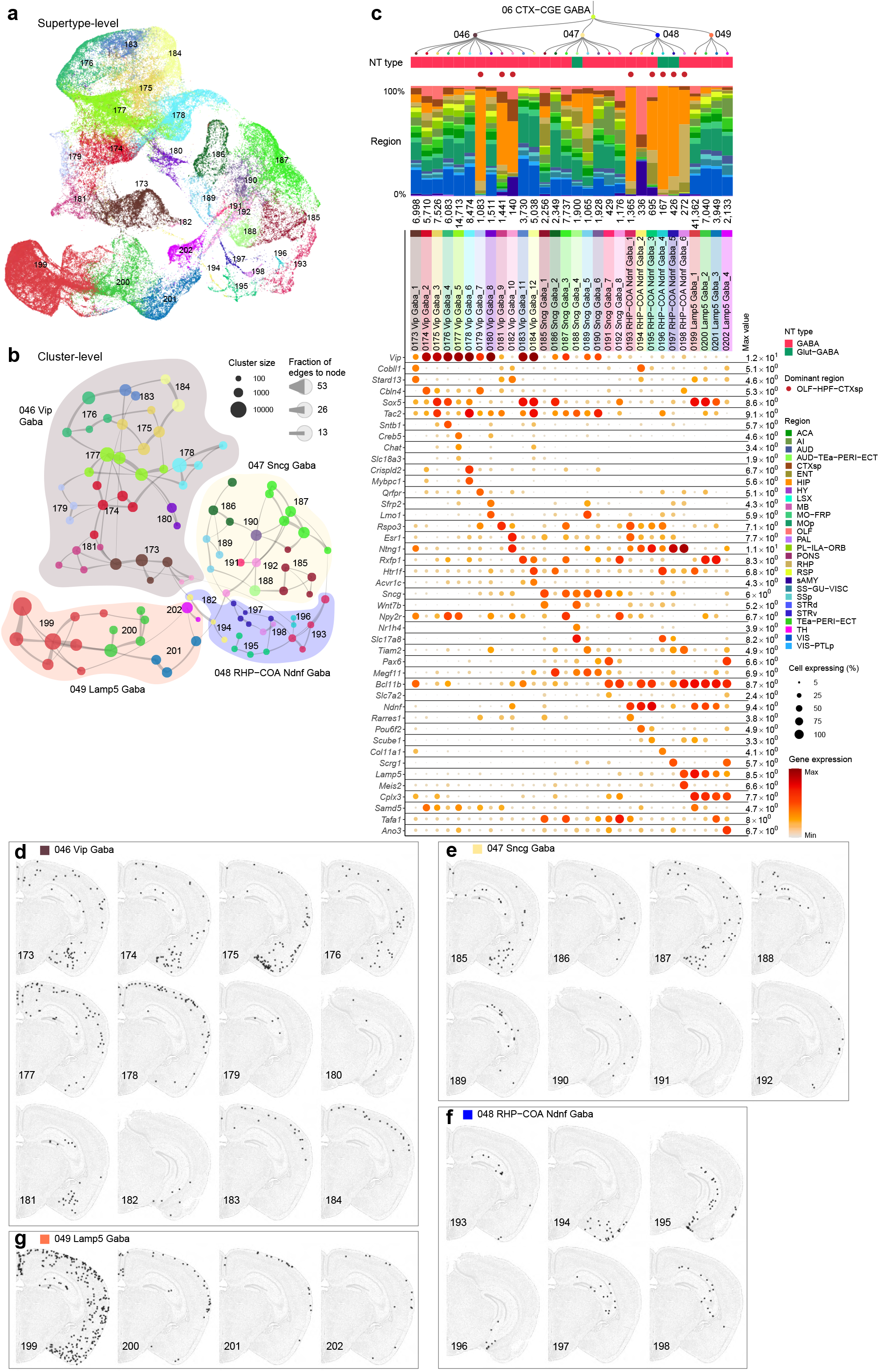
CGE-derived GABAergic neuronal types in the cerebral cortex. **(a)** UMAP representation of all CGE clusters colored by supertype**. (b)** Constellation plot of CGE clusters using UMAP coordinates shown in a. Nodes are colored by supertype and grouped in bubbles by subclass. **(c)** Dendrogram of CGE supertypes followed by bar graphs showing major neurotransmitter type, region distribution of profiled cells, dominant region, and number of cells within supertype, followed by dot plot showing marker gene expression in each supertype from the 10xv3 dataset. The dominant region was assigned if more than 70% of cells are from the assigned region. For the gene expression dot plot, dot size and color indicate proportion of expressing cells and average expression level in each supertype, respectively. **(d-g)** Representative MERFISH sections showing the location of supertypes in CGE subclasses 46 Vip Gaba (d), 47 Sncg Gaba (e), 48 RHP-COA Ndnf Gaba (f), and 49 Lamp5 Gaba (g). Cells are colored and labelled by supertype.

**Extended Data Figure 6.**
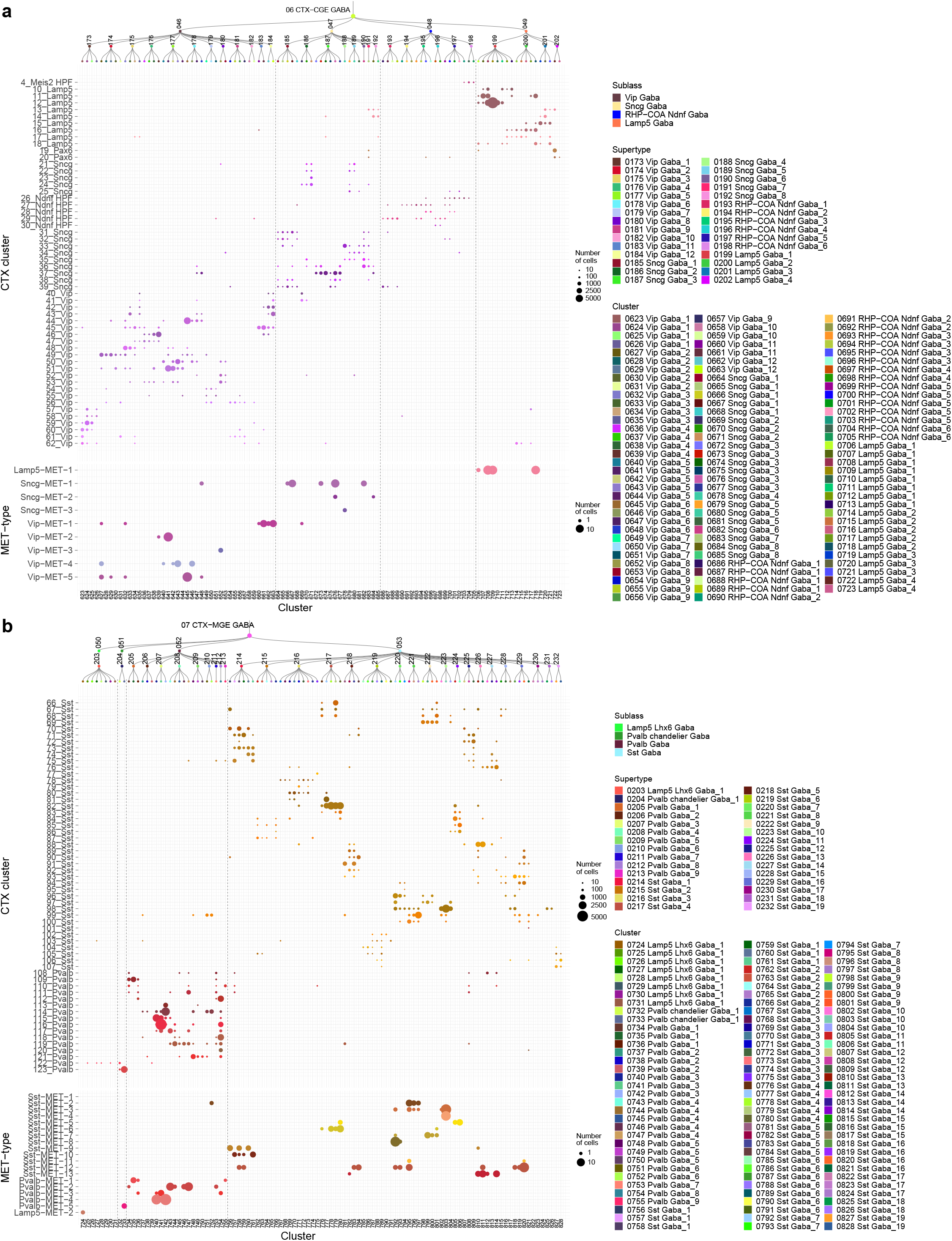
Correspondence of CGE and MGE GABAergic neuronal types with previously published cell-type taxonomies. (**a-b**) CGE (a) and MGE (b) GABAergic cell types identified in this study are compared to cell types in CTX-HPF study^26^ and VISp Patch-seq study^27^. Size of the dots corresponds to the number of overlapping cells in corresponding taxonomies. Columns are separated by supertypes, and rows are separated manually based on subclass in corresponding dataset.

**Extended Data Figure 7.**
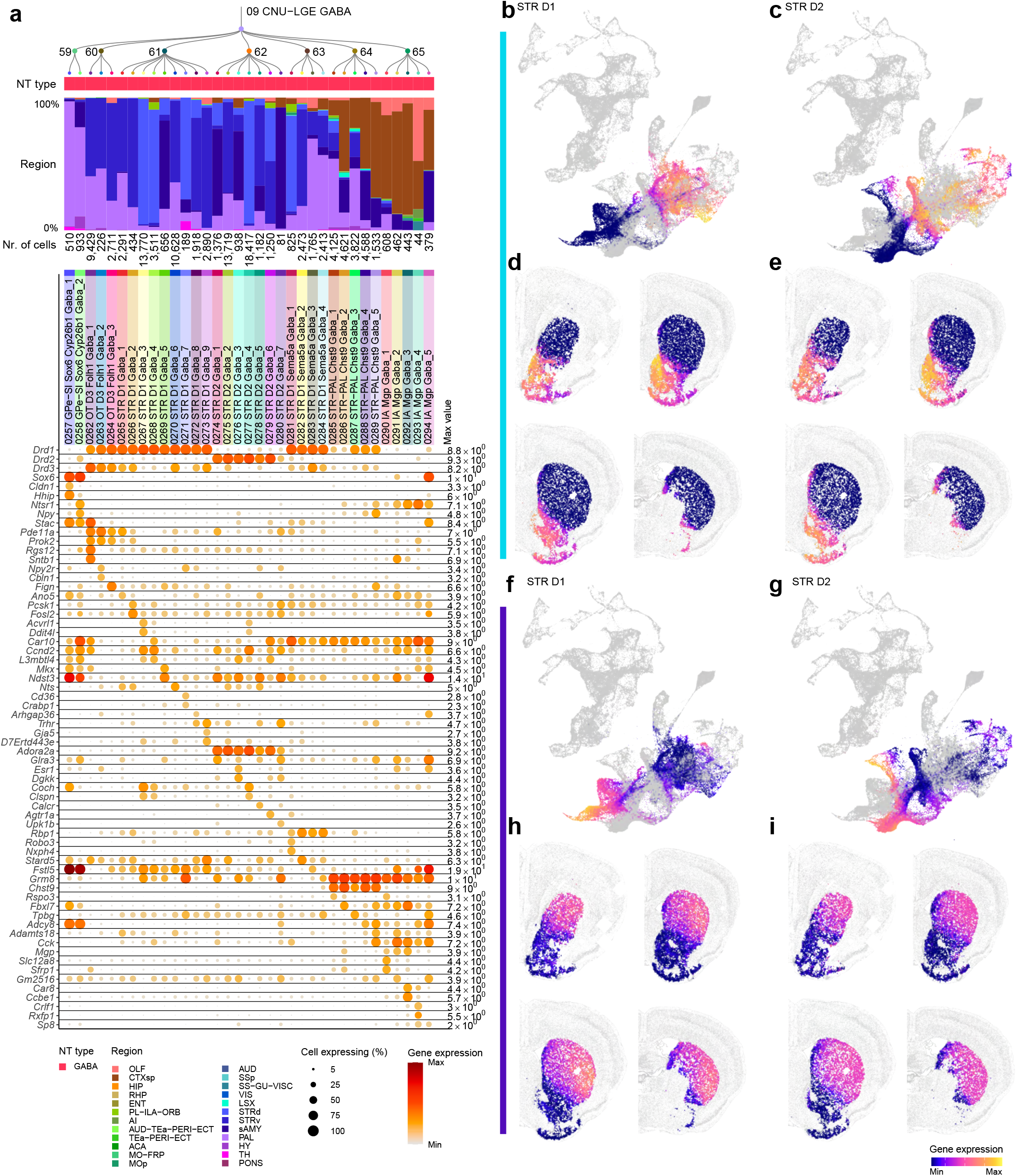
LGE-derived GABAergic cell types of the cerebral nuclei. **(a)** Dendrogram of CNU-LGE supertypes followed by bar graphs showing major neurotransmitter type and region distribution of profiled cells, followed by dot plot showing marker gene expression in each supertype from the 10xv3 dataset. Dot size and color indicate proportion of expressing cells and average expression level in each supertype, respectively. **(b-i)** Based on the gene modules (blue and purple) identified in Figure 5g-h, a cumulative gene score was calculated using UCell. UMAPs showing CNU-LGE GABAergic neurons (b,c,f,g), and representative MERFISH sections (d,e,h,i) colored by blue gene module score for STR D1 (b,d) and STR D2 (c,e) types or colored by purple gene module score for STR D1 (f,h) and STR D2 (g,i) types.

**Extended Data Figure 8.**
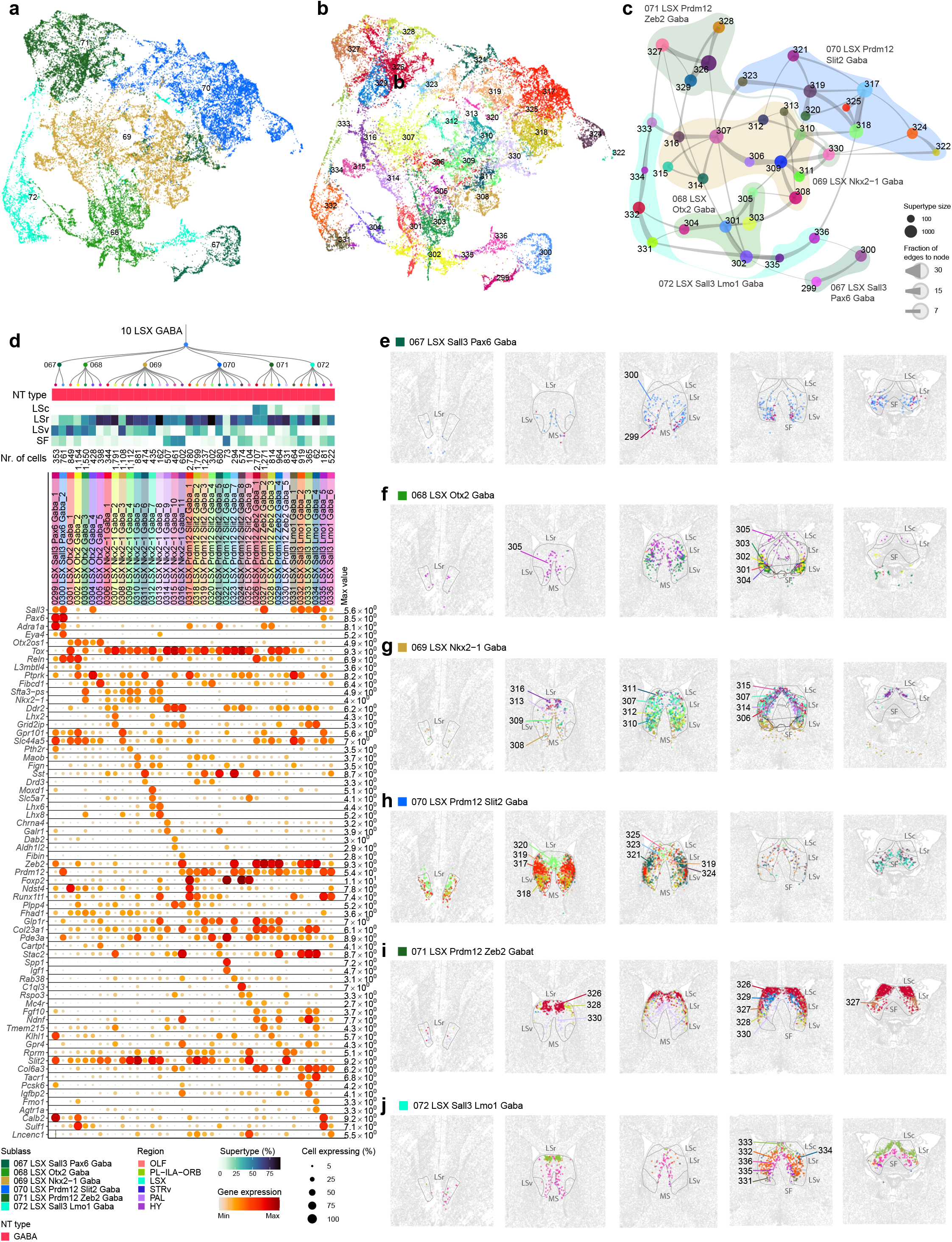
GABAergic cell types of the lateral septum. **(a-b)** UMAP representation of all LSX clusters colored by subclass **(a)** or supertype **(b)**. **(c)** Constellation plot of LSX clusters using UMAP coordinates shown in b. Nodes are colored by supertype and grouped in bubbles by subclass. **(d)** Dendrogram of LSX supertypes followed by tiles showing major neurotransmitter type, followed by a heatmap showing the region distribution of profiled cells, then followed by dot plot showing marker gene expression in each supertype from the 10xv3 dataset. Dot size and color indicate proportion of expressing cells and average expression level in each supertype, respectively. **(e-j)** Representative MERFISH sections showing the location of the LSX subclasses 67 LSX Sall3 Pax6 Gaba (e), 68 LSX Otx2 Gaba (f), 69 LSX Nkx2-1 Gaba (g), 70 LSX Prdm12 ve Gaba (h), 71 LSX Prdm12 do Gaba (i), and 72 LSX Sall3 Lmo1 Gaba (j).

**Extended Data Figure 9.**
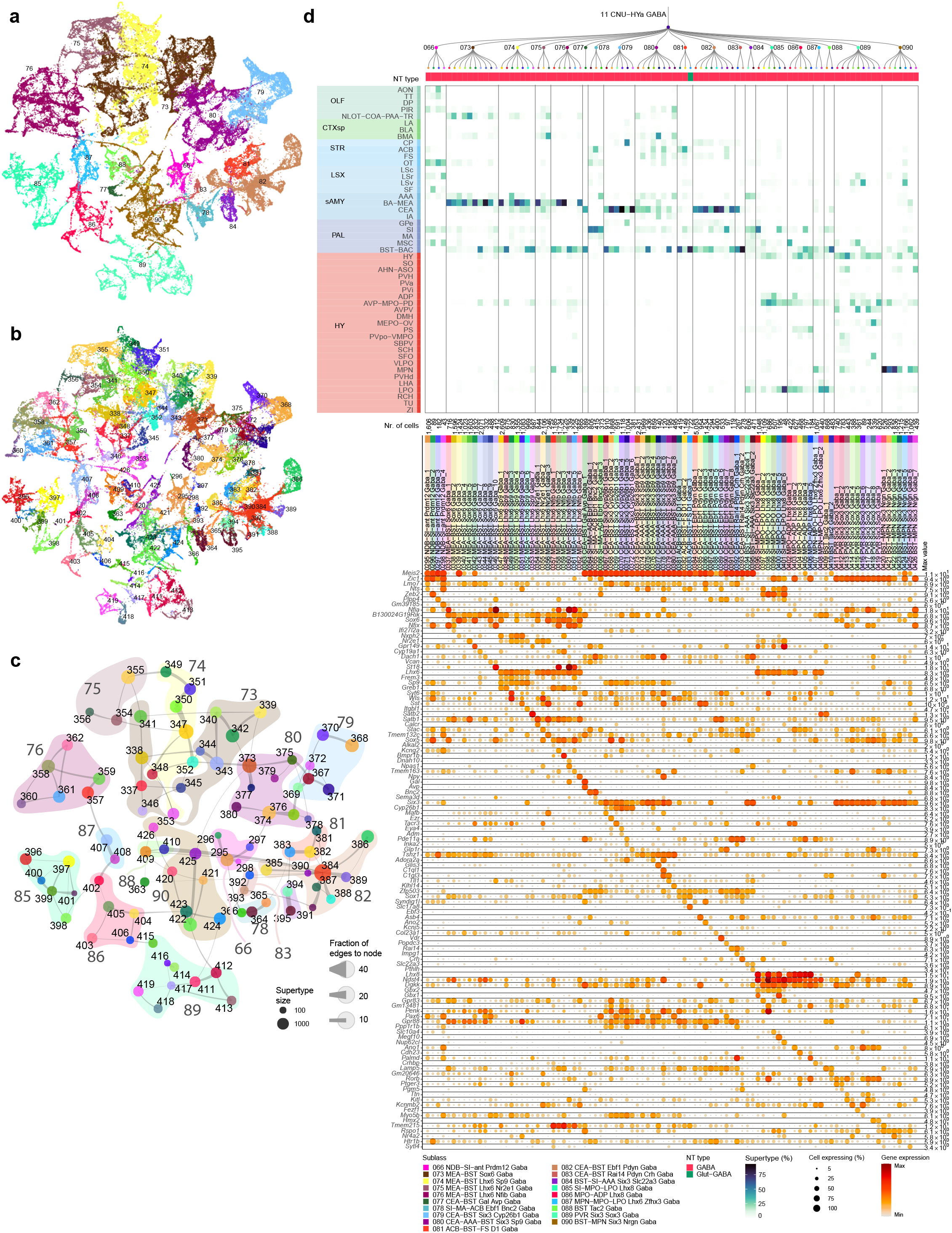
GABAergic cell types of the CNU and anterior hypothalamus (HYa). **(a-b)** UMAP representation of all CNU-HYa clusters colored by subclass (a) or supertype (b). **(c)** Constellation plot of CNU-HYa supertypes using UMAP coordinates shown in b. Nodes are colored by supertype and grouped in bubbles by subclass. **(d)** Dendrogram of CNU-HYa supertypes followed by tiles showing major neurotransmitter type, followed by a heatmap showing the region distribution of profiled cells, then followed by dot plot showing marker gene expression in each supertype from the 10xv3 dataset. Dot size and color indicate proportion of expressing cells and average expression level in each supertype, respectively.

**Extended Data Figure 10.**
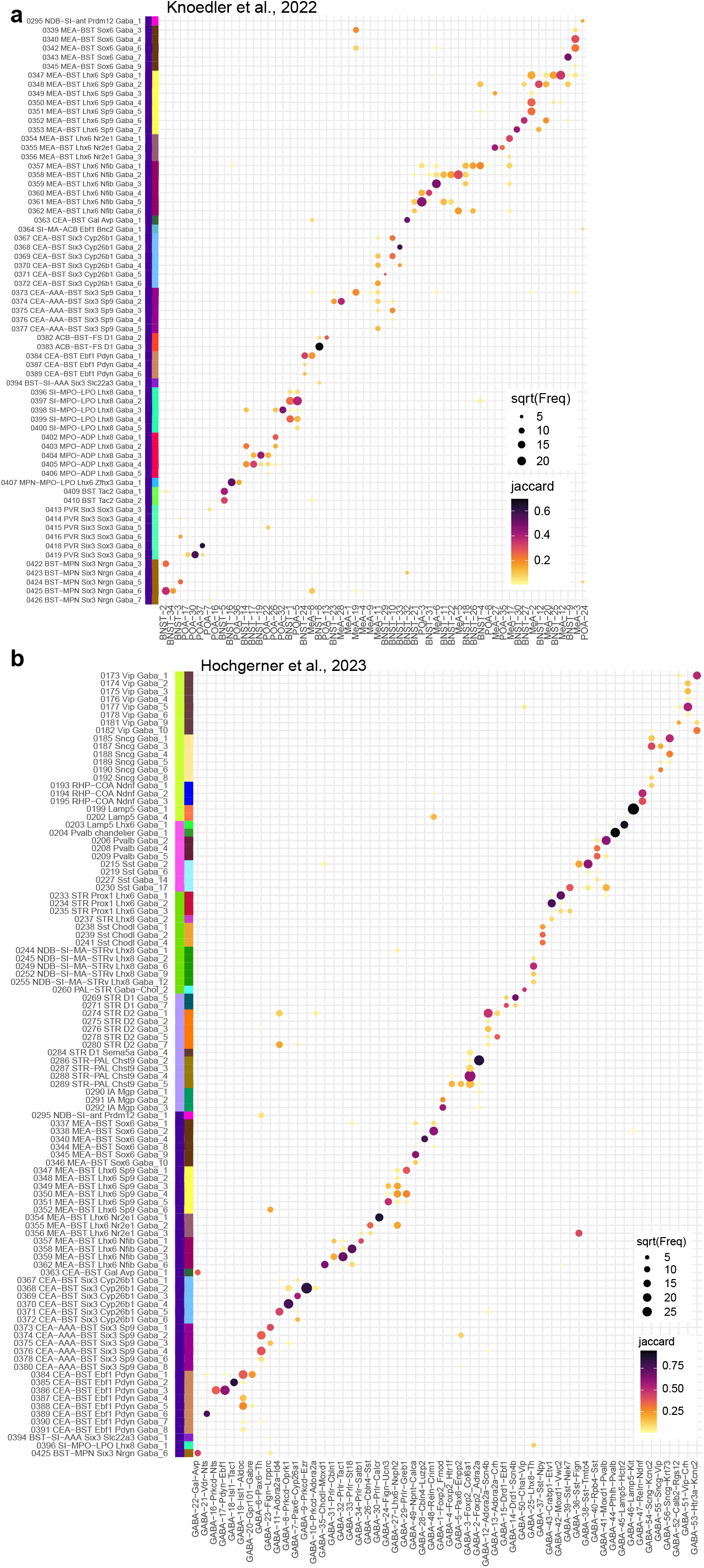
Correspondence between the current transcriptomic taxonomy of the CNU-HYa class and previously published ones. Correspondence was determined by mapping cells from previously published datasets to the current taxonomy as described before. **(a-b)** Mapping of Knoedler et al., 2022 (a) and Hochgerner et al., 2023 (b) to supertypes in the CNU-HYa class or the entire telencephalic GABAergic taxonomy.

**Extended Data Figure 11.**
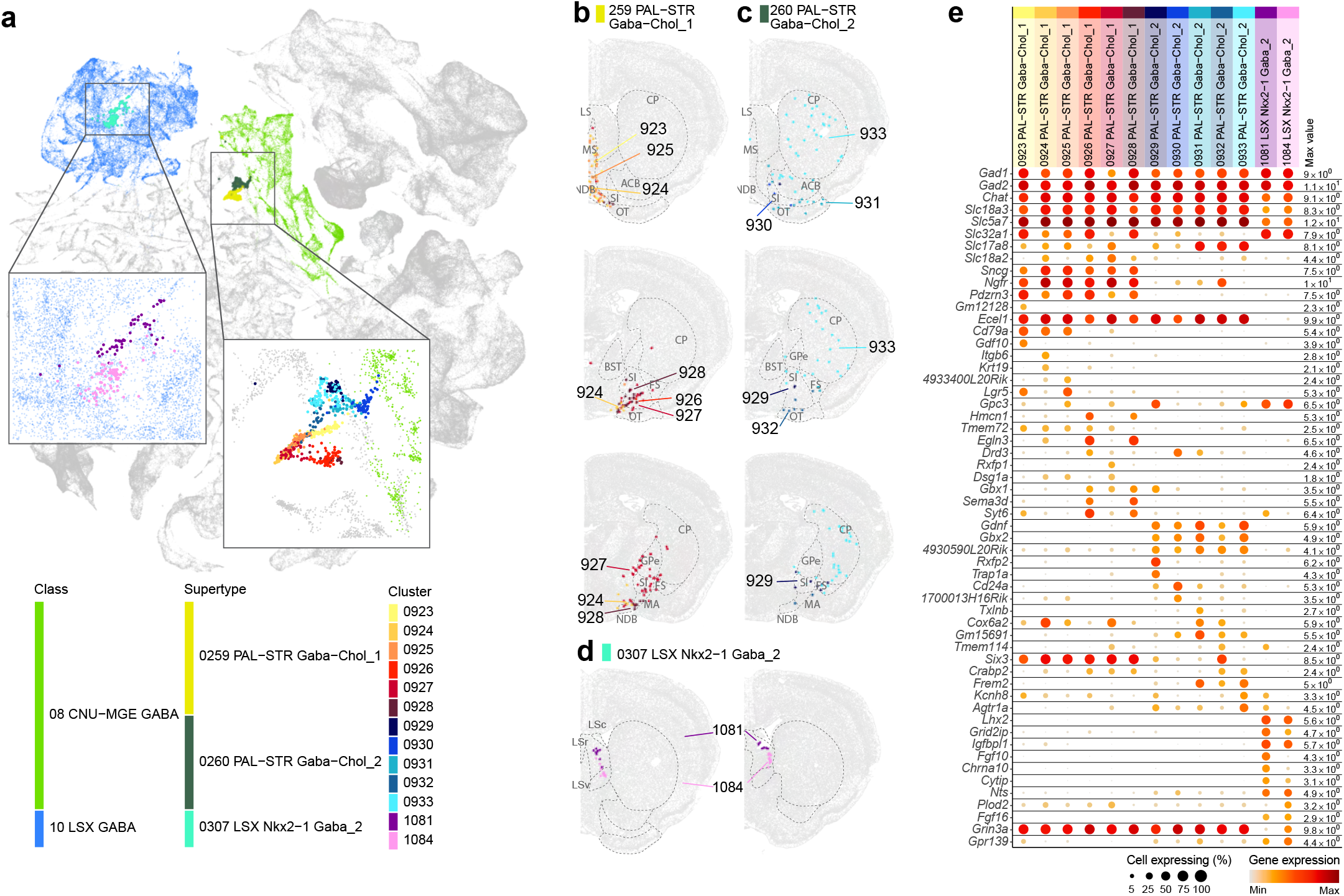
Distribution of basal forebrain cholinergic neurons. **(a)** UMAP representation of all telencephalic GABAergic neurons as in Figure 1b-d, cells in background are colored classes that contain cholinergic neurons and foreground cells are colored by supertype. The insert shows the cholinergic neurons belonging to the CNU GABA class colored by cluster. **(b-d)** Representative MERFISH sections showing cholinergic neurons colored by cluster for supertypes 259 PAL-STR Gaba-Chol_1 (b), 260 PAL-STR Gaba-Chol_2 (c), and 307 LSX Nkx2-1 Gaba_2 (d). **(e)** Dot plot showing marker gene expression in each cholinergic cluster from the 10xv3 dataset. Dot size and color indicate proportion of expressing cells and average expression level in each cluster, respectively.

**Extended Data Figure 12.**
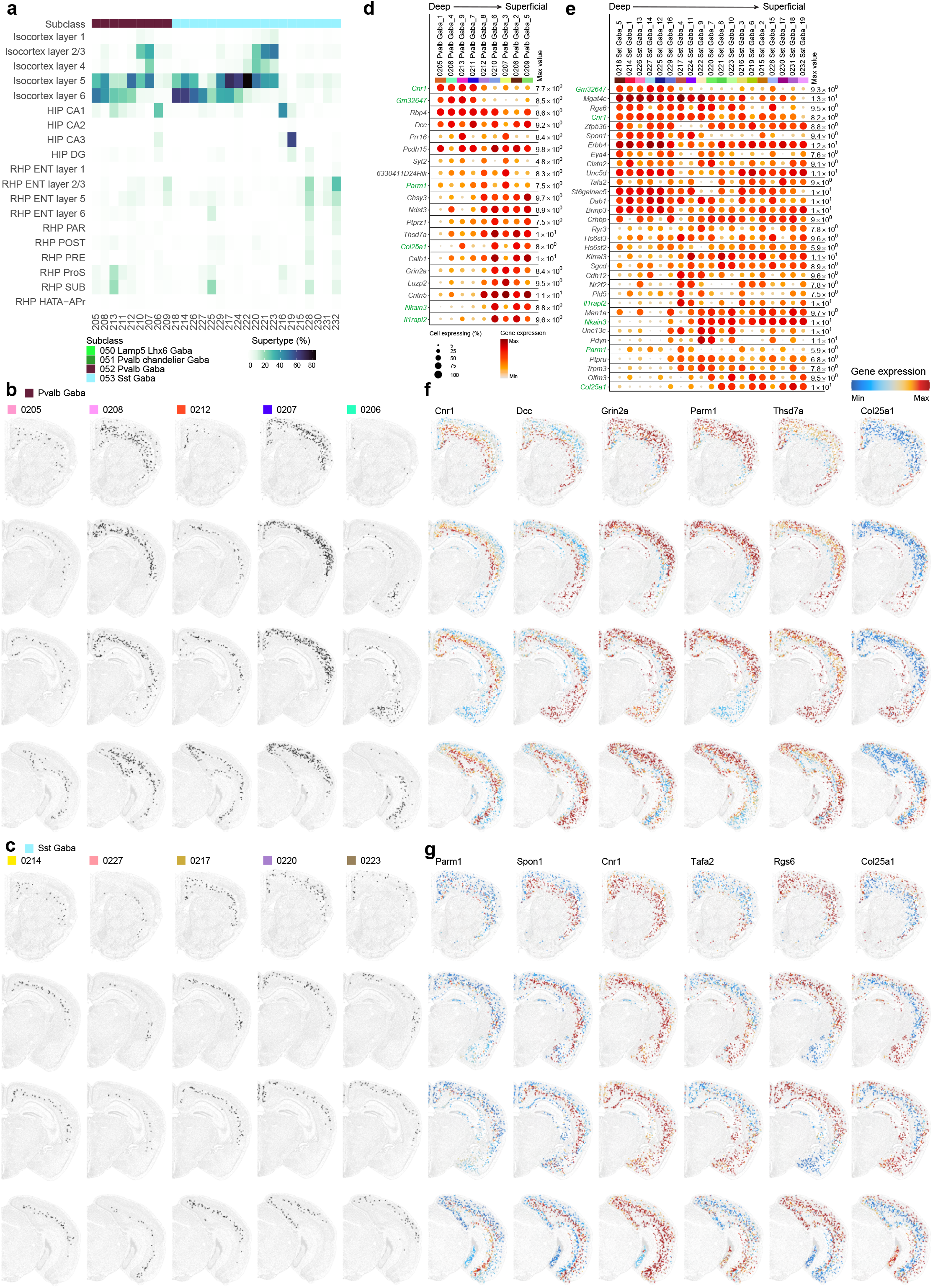
Laminar distribution of MGE GABAergic neurons in cortex and hippocampal formation. (**a**) Heatmap showing the proportion of cells in each layer or region of the isocortex and HPF for supertypes in the MGE-CTX GABA class. (**b,c**) Representative MERFISH sections showing the distribution of neurons across cortex and hippocampal formation in select supertypes from subclasses 42 Pvalb Gaba (b) and 43 Sst Gaba (c). (**d,e**) Dot plot showing expression level of genes driving the gene expression gradient along the cortical depth for supertypes (ordered from superficial to deep) within the Pval Gaba (d) and Sst Gaba (e) subclasses. Dot size and color indicate proportion of expressing cells and average expression level in each supertype, respectively. (**f, g**) Representative examples of genes that drive the laminar gene expression gradient, plotted on the same MERFISH section as in panels b and c, shown for the 42 Pvalb Gaba (f) and 43 Sst Gaba (g) subclasses.

**Extended Data Figure 13.**
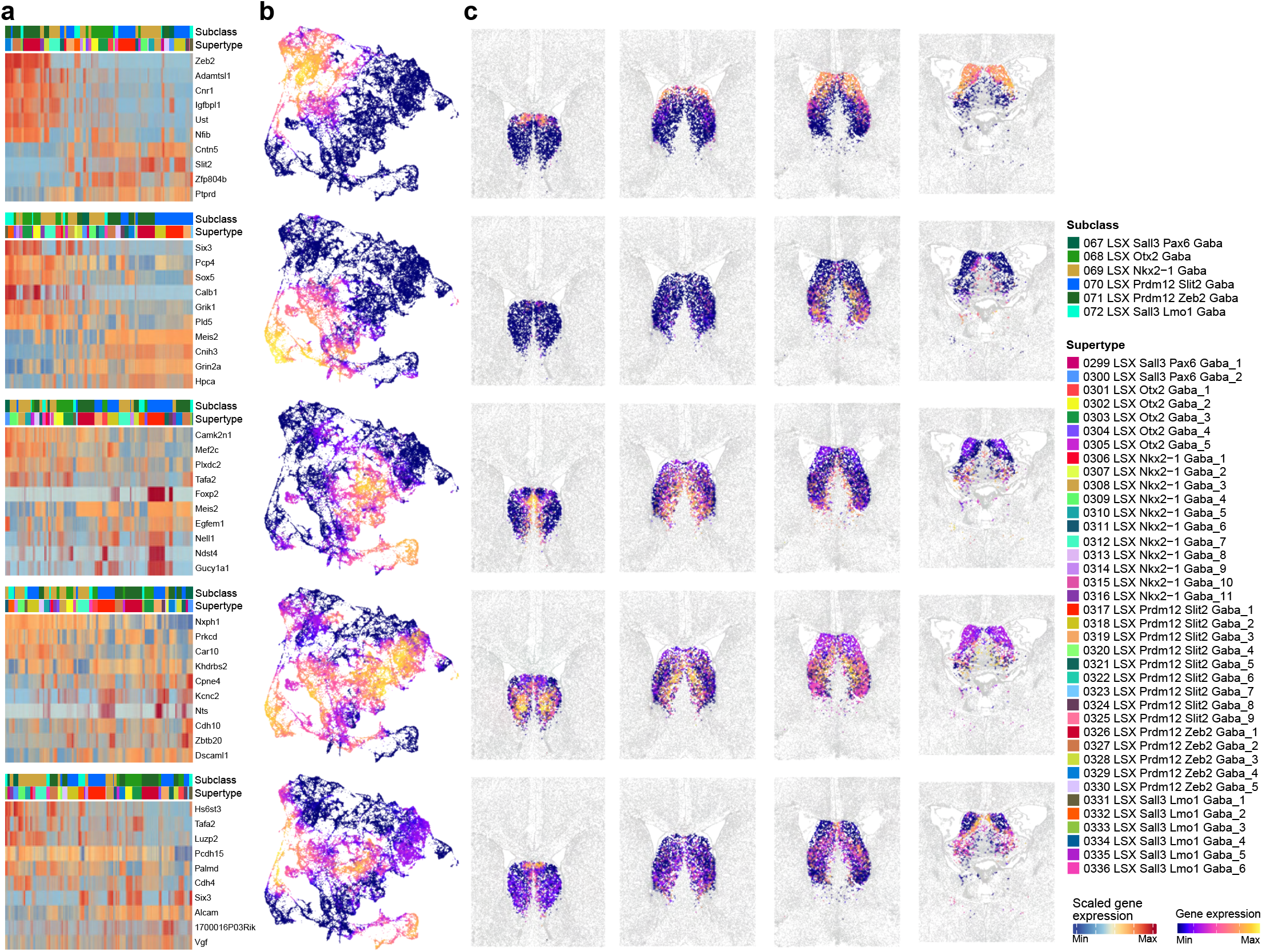
Spatial gradients in LSX. **(a)** Heatmap showing expression of genes that drive the spatial gradients among subclasses in LSX. Top five gene modules containing genes that are both up- and down-regulated along the spatial gradients. We calculated the gene signature score for these modules for every cell and colored the scRNA-seq UMAP representation (**b**) and representative MERFISH sections (**c**) by gene score.

**Extended Data Figure 14.**
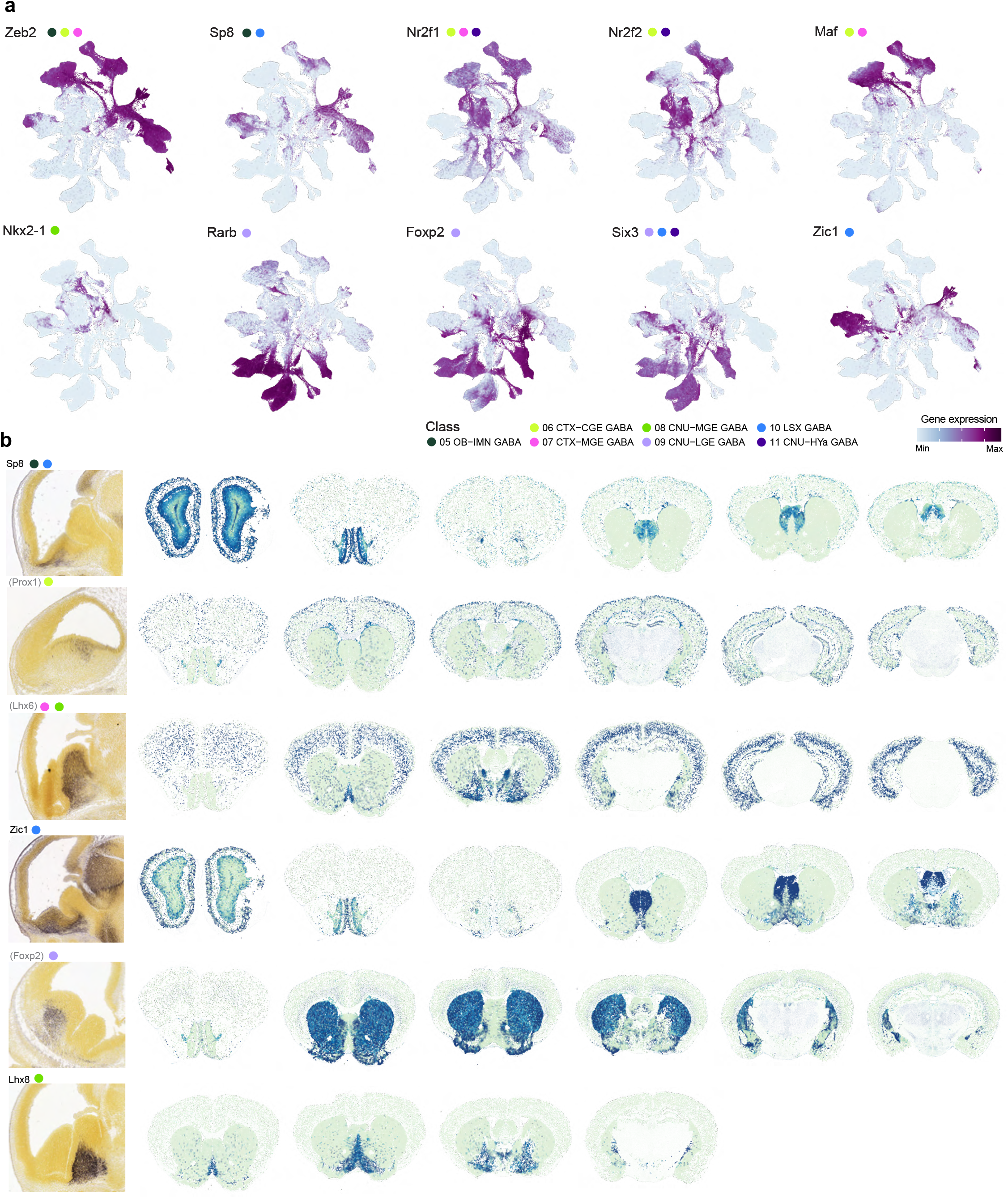
Transcription factor expression marking developmental lineages. **(a)** UMAP representation of all cells across E11-E14, P0, P14 and P56 time points colored by expression level of major lineage marker genes. The colored dots next to each gene name show the classes that express that gene. **(b)** Representative MERFISH sections showing expression of key transcription factors in GABAergic neurons. Genes marked with parentheses show expected spatial gene expression pattern based on imputed data (see **Methods**).

**Extended Data Figure 15.**
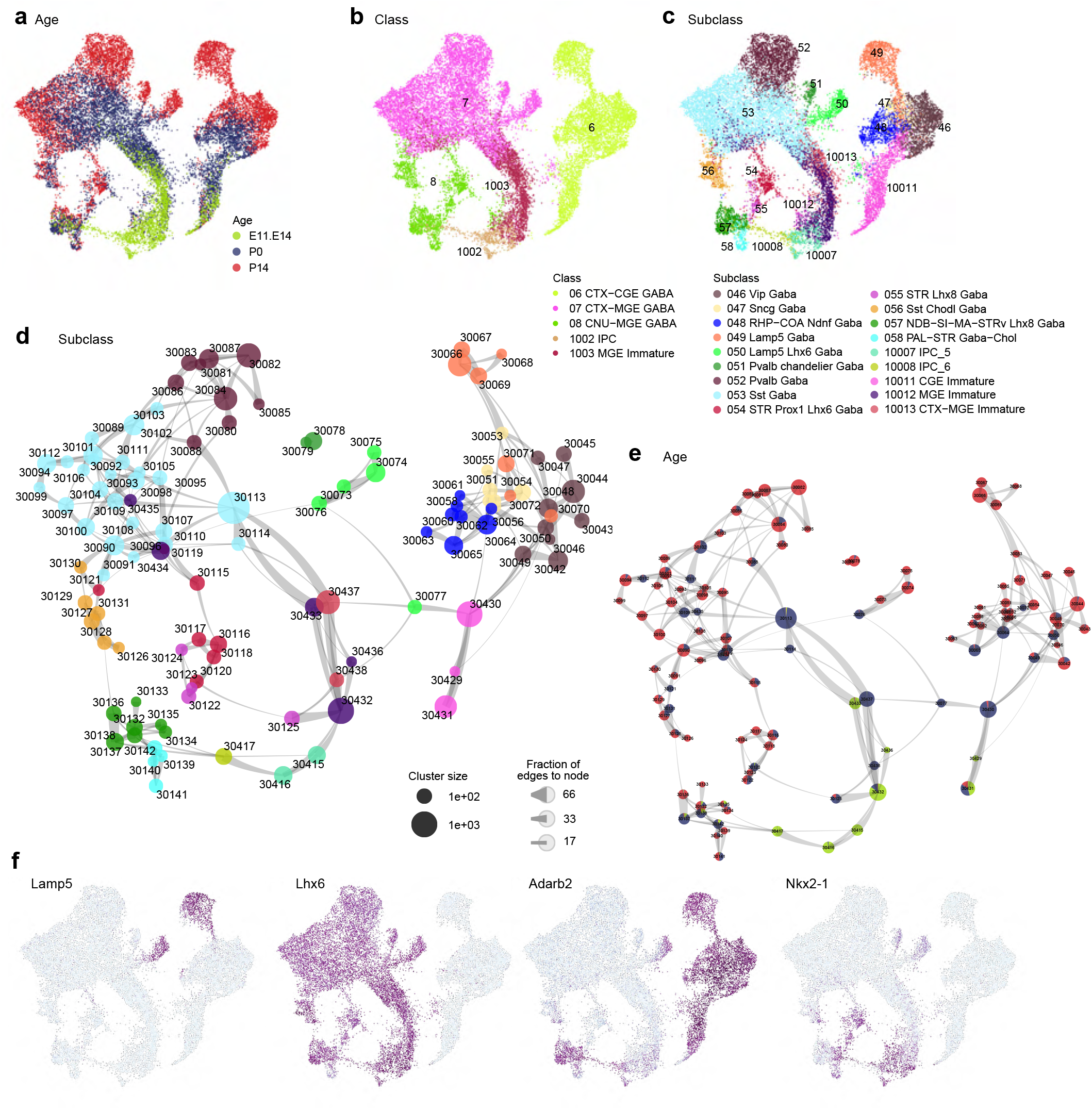
Developmental trajectory of CGE- and MGE-derived neurons. **(a-c)** UMAP representation of all neurons at E11.5-E14.5, P0, and P14 that will form the 06 CTX-CGE GABA, 07 CTX-MGE GABA, and 08 CNU-MGE GABA classes colored by age (a), class (b), or subclass (c). **(d,e)** Constellation plots showing all clusters using UMAP coordinates. Nodes are colored by subclass (d), or proportion of age group (e). **(f)** UMAPs showing expression of genes that link the Lamp5 Lhx6 subclass to both CGE and MGE origins.

**Extended Data Figure 16.**
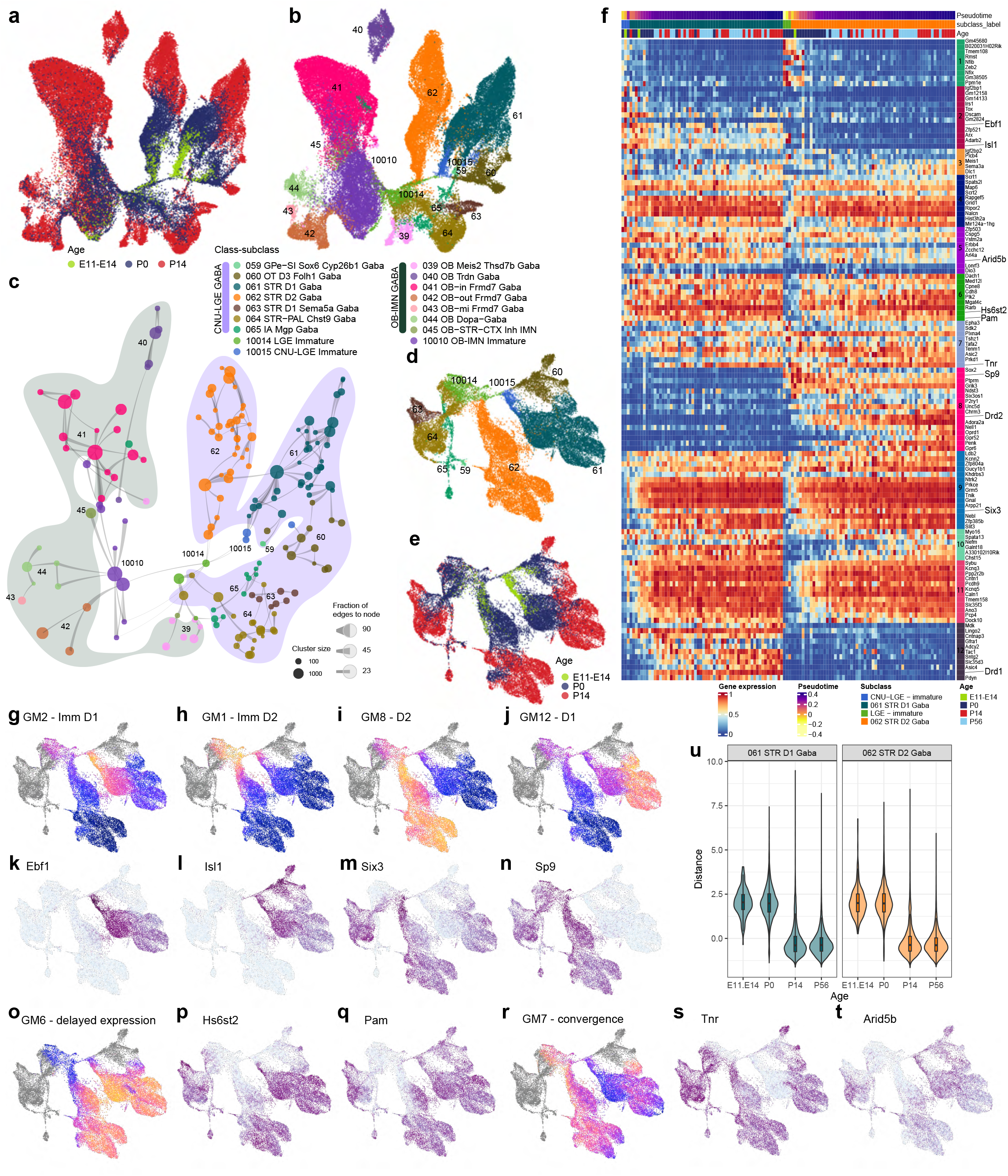
Developmental trajectory of CNU LGE-derived neurons. **(a-b)** UMAP representation of all neurons at E11.5-E14.5, P0, and P14 that are derived from LGE and will populate the OB-IMN and CNU-LGE classes. UMAPs are colored by age (a) or subclass (b). **(c)** Constellation plot showing all developmental clusters using UMAP coordinates from panel a. Nodes are colored by supertype and bubbles behind constellation are colored by subclass. **(d-e)** UMAP representation of all neurons at E11.5-E14.5, P0, and P14 that will form the CNU-LGE class. UMAPs are colored by subclass (d) or age (e). **(f)** Heatmap showing differentially expressed genes in the STR D1 and STR D2 lineages across time. Twelve gene modules were identified that show various modes of expression along and between subclasses over time. (**g-j**) Gene module scores marking different stages along the maturation path of D1 and D2 neurons. Gene modules 2 and 1 highlight immature STR D1 (g) and immature STR D2 (h) neurons respectively, while gene modules 12 and 8 mark mature STR D1 (i) and mature STR D2 (j) neurons respectively. (k-n) UMAP representation like in panels d-e colored by major lineage markers. (**o-q**) Gene module 6 contains genes highlighting the delayed maturation of STR D2 vs STR D1 neurons (o), such as two exemplar genes *Hs6st2* (p) and *Pam* (q). **(r-t)** Gene module 7 contains genes whose expressions converge along the maturation trajectory of STR D1 and STR D2 neurons (r), such as two exemplar genes *Tnr* (s) and *Arid5b* (t). **(u)** Violin plot showing the transcriptomic distance between STR D1 and STR D2 subclass transcriptomes across the time course.

**Extended Data Figure 17.**
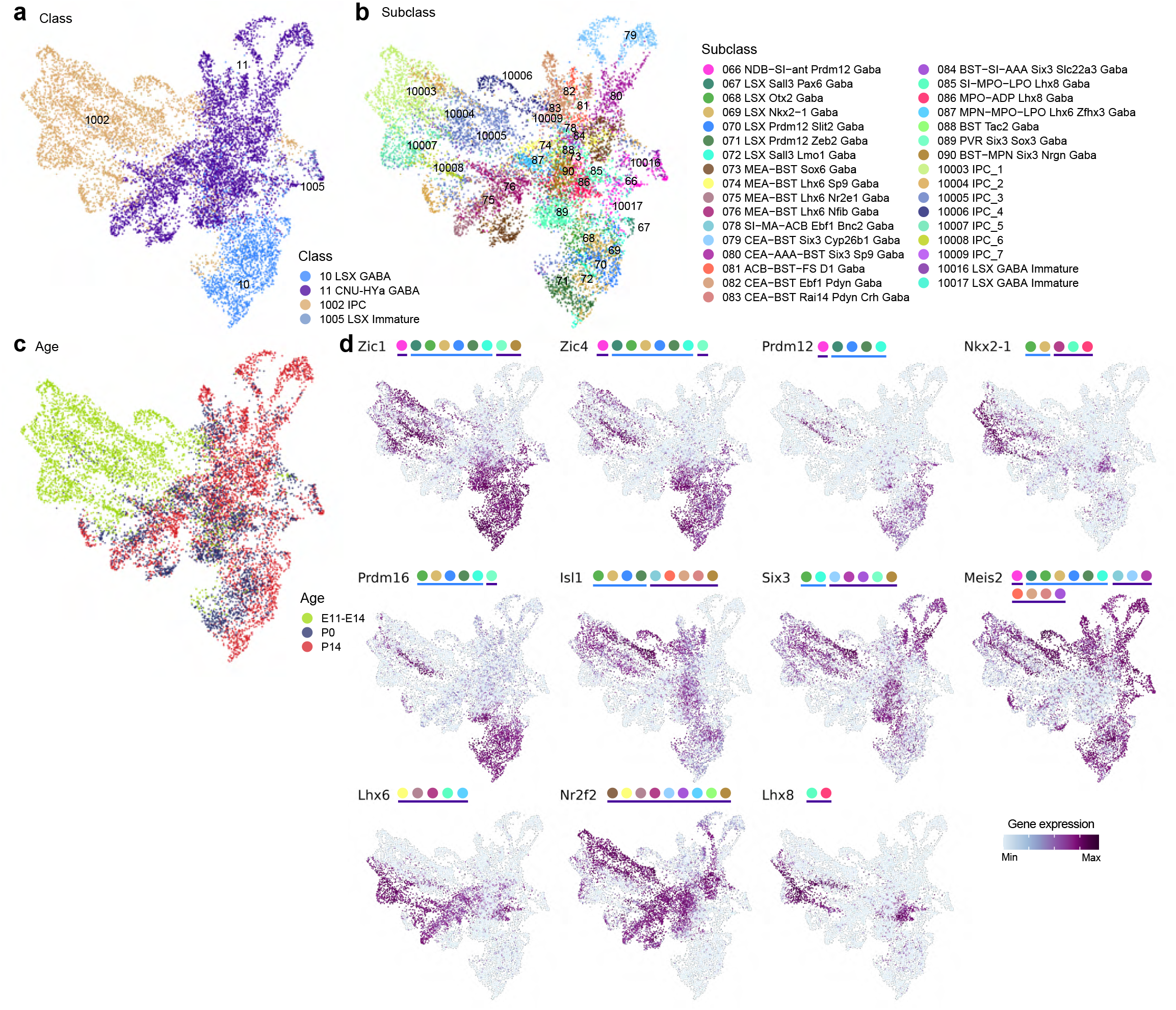
Developmental trajectory of LSX and CNU-HYa GABAergic neurons. **(a-c)** UMAP representation of all neurons at E11.5-E14.5, P0, and P14 that will form the 10 LSX GABA and 11 CNU-HYa GABA classes colored by class (a), subclass (b), and age (c). **(d)** UMAP representation as in panels a-c colored by expression level of subclass marker genes. The colored dots next to each gene name show the subclasses expressing that gene.

**Supplementary Table 1. Allen Mouse Brain Common Coordinate Framework version 3 (CCFv3) regional ontology.** Regions outside telencephalon are greyed out. Adopted from Wang et al, 2020.

**Supplementary Table 2. MERFISH 500-gene panel used in Vizgen MERSCOPE platform to generate the whole mouse brain MERFISH dataset.**

**Supplementary Table 3. Cell type taxonomy of GABAergic neuronal types in the telencephalon.** This taxonomy was defined as the Subpallium-GABA neighborhood in the whole mouse brain cell type atlas in Yao et al, 2023. Detailed information for telencephalic GABAergic clusters, including membership at different levels (supertype, subclass, class, and division), NT type, NT type combo, major NT marker genes, major neuropeptides, top and combo marker genes, main dissection region, manual anatomical annotation, number of 10x v2 and 10x v3 cells, fraction of male and female cells, and accession numbers to cell types.

**Supplementary Table 4. Developmental cell type taxonomy of GABAergic neuronal types in the telencephalon.** Detailed information for developmental GABAergic clusters, including membership at different levels (supertype, subclass, and class), number of cells from each age group, and top marker genes.

**Supplementary Table 5. Donor information of the Developmental scRNA-seq dataset.** All donors used to generate the developmental scRNA-seq data in this study are listed, with associated metadata including sex, age, genotype, etc. From one donor multiple regions could be dissected.

